# Extracellular and intracellular components of the impedance of neural tissue

**DOI:** 10.1101/2021.08.05.455210

**Authors:** Claude Bedard, Charlotte Piette, Laurent Venance, Alain Destexhe

## Abstract

Electric phenomena in brain tissue can be measured using extracellular potentials, such as the local field potential, or the electro-encephalogram. The interpretation of these signals depend on the electric structure and properties of extracellular media, but the measurements of these electric properties are still debated. Some measurements point to a model where the extracellular medium is purely resistive, and thus parameters such as electric conductivity and permittivity should be independent of frequency. Other measurements point to a pronounced frequency dependence of these parameters, with scaling laws that are consistent with capacitive or diffusive effects. However, these experiments correspond to different preparations, and it is unclear how to correctly compare them. Here, we provide for the first time, impedance measurements (in the 1-10 KHz frequency range) using the same set-up in various preparations, from primary cell cultures to acute brain slices, with a comparison with similar measurements performed in artificial cerebrospinal fluid with no biological material. The measurements show that when the current flows across a cell membrane, the frequency dependence of the macroscopic impedance between intracellular and extracellular electrodes is significant, and cannot be captured by a model with resistive media. Fitting a mean-field model to the data shows that this frequency dependence could be explained by the ionic diffusion mainly associated to Debye layers surrounding the membranes. We conclude that neuronal membranes and their ionic environment induce strong deviations to resistivity, that should be taken into account to correctly interpret extracellular potentials generated by neurons.

**Significance:** The electro-encephalogram recorded at the scalp surface and local-field potentials recorded within neural tissue are generated by electric currents in neurons, and thus depend on the impedance of neural tissue. Different measured values were proposed, and it is currently unclear what is the real impedance of neural tissue. Here, we show that the impedance depends on the measurement technique. If the measurement is exclusively extracellular, the system appears as equivalent to a simple resistor. However, if the measurement includes an intracellular electrode, a more complex impedance is observed, because the current has to flow through the membrane, as happening in the brain. Thus, we provide an explanation for apparent disagreements, and indicate in which cases each impedance should be used.

## 1 Introduction

The genesis of extracellular electric potentials in the brain depends on the electric properties of the extracellular medium. The exact nature of these electric properties is important, because non-resistive media will necessarily impose frequency-filtering properties to electric signals [1, 2] and therefore will influence any source localization. Early studies modeled the genesis of extracellular potentials assuming that the medium is analogous to a resistance [3]. Evidence for such resistive media was provided by different measurements [4, 5, 6], while other measurements [7, 8, 9] revealed a more complex situation, where the measured electric parameters displayed a dependence on frequency in contrast to the frequency-independence of resistive systems. Computational models showed that such a dependence on frequency can be obtained if there are strong spatial variations of conductivity and/or permittivity [10]. Further models showed that frequency-dependent electric parameters can also result from electric polarization of the medium [11], or from ionic diffusion [12].

The extracellular medium properties were also estimated indirectly by correlating intracellular and local field potentials (LFP) [13] or by the frequency-dependence between electro-encephalogram (EEG) and magneto-encephalogram signals (MEG) [14]. Although a framework was proposed to explain the contradictory measurements [15], the electric nature of the extracellular medium is still debated [15, 16, 17].

The main problem to resolve this debate is that different experiments correspond to very different preparations, and it is not clear how to correctly compare them. For example, impedance was either measured intracellularly [9] or extracellularly [5, 6]. Could this account for discrepancies observed across studies ? In the present study, we provide for the first time, impedance measurements, either with an intracellular electrode or extracellularly, in different prepearations, in acute brain slices, in primary cell cultures, and we compare to measurements using the same set-up in artificial cerebrospinal fluid (ACSF).

## 2 Materials and Methods

### 2.1 *In vitro* electrophysiology

#### Animals

C57BL/6 and Swiss mice were housed by groups of 3-5 mice, in a 12h light/dark cycle, with food and water available *ad libitum*. All experiments were performed in accordance with local animal welfare committee (Center for Interdisciplinary Research in Biology, and Institut de Biologie Paris-Seine, IBPS, Ethical Committees) and EU guidelines (Directive 2010/63/EU). Every precaution was taken to minimize stress and the number of animals used in each series of experiments.

#### Neuronal primary cultures

The brains (from Swiss mice) were removed from day 14 embryos and striata were isolated and dissociated by gently pipetting in PBS-0.6% glucose. Cells were collected by centrifugation at 1000 x g for 5 min. Cell pellets were resuspended in Neurobasal medium supplemented with B27 (Invitrogen, Thermo Fisher Scientific, Illkirch, France), 500 nM L-glutamine, 60 *μ*g/ml penicillin-streptomycin and 25 *μ*M *β*-mercaptoethanol (Sigma, Saint-Quentin Fallavier, France) and then plated into 24-well (1.8 x 10 ^5^ cells per well) plates coated with 50 *μ*g/ml poly-d-lysine (Sigma). After removal of the coating solution, cells were seeded in the Neurobasal medium on glass cover-slips and cultured at 37°C in 95% air and 5%*CO*_2_. When placed in the recording chamber, the cell culture was initially superfused with a 95% *O*_2_ /5 % *CO*_2_-bubbled Neurobasal medium, and then progressively diluted in ACSF solution. Patch-clamp recordings were made from day 3 to day 10 after seeding.

#### Brain slices preparation

Horizontal brain slices (from C57BL/6) with a thickness of 300 *μ*m were prepared from postnatal P30-40 mice using a vibrating blade microtome (VT1200S; Leica Biosystems, Nussloch, Germany). Brains were sliced in a 95%*O*_2_ / 5%*CO*_2_-bubbled, ice-cold cutting ACSF solution containing: *NaCl* 125 *KCl* 2.5 mM, glucose 25 mM, *NaHCO*_3_ 25 mM, *NaH*_2_*PO*_4_ 1.25 mM, *CaCl*_2_ 1 mM, *MgCl*_2_ 1 mM, and pyruvic acid 1 mM and then transferred into the same solution at 34°C for one hour before cell recording.

#### Whole-cell patch-clamp recordings and experimental set-up

Electrophysiological recordings were performed in the dorsal striatum, which has the advantage of having no laminar organization (with neurons presenting a relatively constricted dendritic arbor with a spherical distribution), which hence limits the influence on the current trajectories and avoid strong anisotropic equipotential surfaces. Patch-clamp recordings were combined with extracellular recording using a 2-3.5 MΩ patch-clamp glass pipette. The latter was located within a close vicinity (≈ 5-10 *μ*m) of the patched neuron (Fig. 1A). In the primary cell culture experiments, the average distance separating neighbouring cells was on average 17.4 *μ*m (± 7.1, n=14) and 12.8 *μ*m (± 6.4, n=25) for immature and mature cells respectively, thus about three times larger than the distance separating the two pipettes. Borosilicate glass pipettes of 5–7 MΩ impedance contained for whole-cell recordings: K-gluconate 122 mM, *KCl* 13 mM, HEPES 10 mM, phosphocreatine 10 mM, ATP-Mg 4 mM, GTP-Na 0.3 mM, and EGTA 0.3 mM (adjusted to pH 7.35 with *KOH*). The composition of the extracellular solution and inside the extracellular pipette was the same ACSF solution that was used for brain slices incubation. Signals were amplified using EPC9-2 amplifiers (HEKA Elektronik, Lambrecht, Germany) with a very high input impedance (1 TΩ) to ensure there was no appreciable signal distortion imposed by the high impedance electrode [18, 19]. All recordings were performed at 34°C using a temperature control system (Bath-Controller V; Luigs & Neumann, Ratingen, Germany) and brain slices or primary cell cultures were continuously superfused at 2 ml/min with the extracellular solution. The extracellular solution used in the recording chamber had the same ionic composition for all the experimental conditions. Neurons were visualized on a BX51WI microscope (Olympus, Rungis, France) using a 40x/0.80 water-immersion objective for localizing cells for whole-cell recordings and extracellular electrode positioning. Series resistance was not compensated. Current-clamp recordings were sampled at 50 kHz using the Patchmaster v2×73 program (HEKA Elektronik).

**Figure 1:**
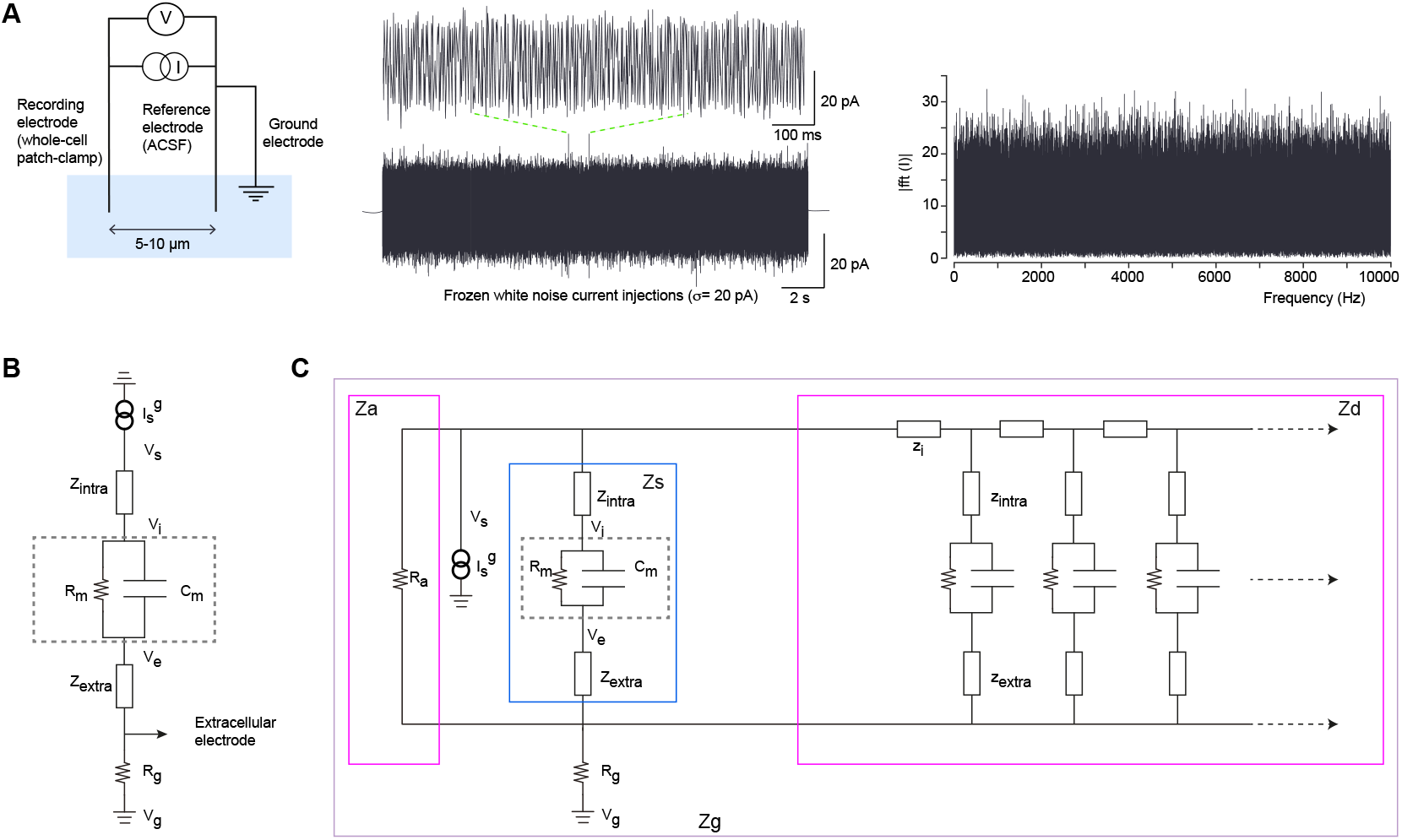
Method used for macroscopic impedance measurements and numerical simulations in different preparations. **(A)** Scheme of the experimental set-up. *Left*: Two borosilicate micropipettes with silver-chloride electrodes and a ground electrode are placed in the ACSF solution. One electrode is used in whole-cell configuration to send white-noise current and simultaneously measure the voltage response of the recorded neuron (acquisition sampling rate: 50 kHz); the other electrode used as a reference is located 5-10 *μ*m away from the recorded neuron. *Middle*: Frozen Gaussian white noise (amplitude: ± 5 to 20 pA; duration: 20 sec) is injected repeatedly into patched neurons. *Right*: Fourier modulus of the injected current as a function of frequency confirms that each frequency from 0 to 10 000 Hz is uniformly sampled. **(B)** Equivalent electrical circuit between intracellular and extracellular electrodes. **(C)** Equivalent electrical circuit between the extracellular electrode and the ground. *Z_a_* is the impedance of the region defined by the isopotential surface 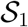 passing through the extracellular electrode and the first isopotential surface 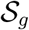 that totally includes the neuron, *Z_s_* is the impedance of the part of the soma included between 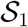 and 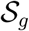, *Z_d_* is the input impedance of the dendrite between surfaces 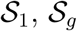, and *Z_g_* is the impedance of the region between 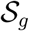 and the ground. *Z_g_* = (*Z_a_* || *Z_s_* || *Z_d_*) ⊕ *R_g_* where || means “in parallel” and ⊕ “in series”.

#### White noise stimulation protocols and signal analyses

Frozen white noise stimuli were applied via the recording patch-clamp electrode in current-clamp mode. A quantity of 20 seconds of Gaussian white noise with zero mean and 5, 10 or 20 pA variance was injected. For each cell, we injected up to 71 times the same sequence of white noise (Fig. 1A). For each neuron, we computed the IV-curve by applying hyperpolarizing current steps of different intensities in current-clamp. Neurons for which the recording voltage of the responses was not inside the linear region of the IV-curve were excluded (only two neurons from brain slices had to be excluded for this reason in this study). For each trial, we calculated as a function of the frequency the modulus and Fourier phase of the voltage difference between the intracellular recording and extracellular reference electrode as well as between the extracellular reference and ground electrode. We then averaged these measures to obtain a Fourier spectrum, ranging from 1-10 kHz, for each cell. Going at higher frequencies was challenging because of the limitations in sampling frequency of our electrophysiological set-up and because the power of the signal becomes weak and dominated by instrumental noise. In addition, the 10 kHz upper limit was sufficient to observe the capacitive effect between the extracellular and intracellular electrodes, because of their close proximity.

Importantly, the numerical measurement of the phase induces a small delay Δ*t* between the real and digitally measured voltage, and this delay depends on the sampling frequency. This delay is likely due to the electronics and does not affect the modulus of the impedance, but is proportional to frequency. We have measured and corrected for this effect using the measurement of the resistance between the two electrodes in ACSF. Without this correction, we would have Φ(*ν*) = 0 because saline is resistive for *v* between 0 and 10 kHz. With the delay, we have 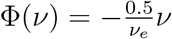 where *ν_e_* is the sampling frequency.

We have fitted the models on the experimentally-measured phase values such as to minimize the mean square distance between experimental and theoretical values. To do this, we have computed the average values on small intervals of frequency (≈ 1 *Hz*) in the impedance modulus of the experimental data in each example. In other words, we have calculated 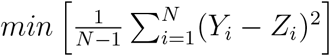, where *Z* and *Y* are respectively the average value over a segment of length equal to 1 Hz in the model and experimental data. We have adjusted the parameters of the different models to minimize the largest absolute distance between the modulus over the ensemble of intervals, between the average experimental and model values. This choice simplifies the comparison between the different models and experimental results. Importantly, log-log graphs give the false impression that there is more noise at high frequencies, but this is due to the high density of points.

Finally, we fitted numerically the polar representation of the experimental data in Fourier frequency space, using a filtering with the cubic spline method [25]. This method has a smooth derivative, which is appropriate for very noisy modulus and phase values. Also, the data were fit to models that are plausible biophysically (resistive, diffusive, etc), so that the parameters have a clear biological or physical interpretation. In all cases, each biophysical model has a few parameters (which number remains very small compared to the data set), and provides us with estimates of these parameters, as we describe in the Results section.

### 2.2 Mathematical and Physical models used to simulate the experimental data

To model the experimental data, we use Maxwell equations under the electric quasistatic approximation^1^ which was formulated in mean-field in previous studies [12, 20]. At the first-order of this mean-field theory, the macroscopic impedance^2^ (in Fourier frequency space) of a point neuron in a heterogeneous medium is given by:

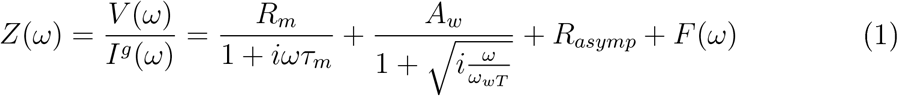

where 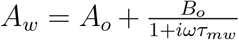. This expression was written by expliciting the following physical phenomena. The first right-hand term corresponds to the usual R-C circuit of the membrane, the second term is due to ionic diffusion (in a linear approximation) as well as to electric polarization (if the polarization relaxation time is negligible). The last term takes into account other physical phenomena which may introduce a frequency dependence in the extracellular medium (such as capacitive effects^3^). The physical meaning of the parameters is as follows: *R_m_* is the macroscopic membrane resistance, *τ_m_* = *R_m_C_m_* is the membrane time constant, *A_ω_* is the amplitude of the diffusive impedance (which is a real number when the polarization relaxation time (Maxwell-Wagner time *τ_MW_*) is negligible, *R_asymp_* is the asymptotic resistance for very high frequencies, and *ν_wT_* = *ω_wT_*/2*π* is its Warburg threshold frequency (for details, see Supplemental Information, Appendix C).

The model used to take into account the effect of dendrites is the same as used previously [9], and consists of a model with a soma and a dendrite. Because we have a monopolar current source (electrode injection), the electrotonic length of a dendritic branch is given by:

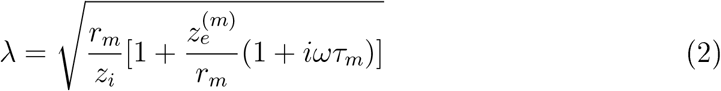

where the following quantities are defined in the generalized cable: *z_i_* is the cytoplasm impedance per unit length, *r_m_* is the specific membrane resistance, and 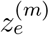 is the specific input impedance of the extracellular medium as seen by the membrane.

We have 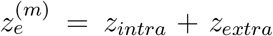 (from Fig. 1). The physical link between the generalized cable is the following. The current density in the dendritic stick has a component in parallel to the axis of the stick, and a component perpendicular to it. However, the electric conductivity is different for these two components. *z_i_* is associated to the parallel component, where the current density is physically related to the cytoplasm, and *z_intra_* is associated to the perpendicular component, which is physically related to the Debye layers in the inner side of the membrane. We have

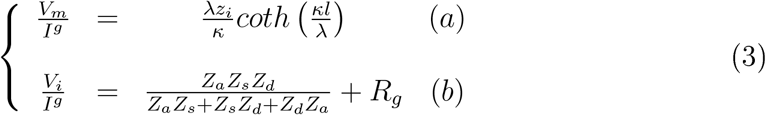

where the neuronal cable model in an “open configuration” (i.e, without return current) was used, as defined in [22].

Because the current that goes from the neuron to the extracellular space separates into several parts, one part that flows through the soma membrane, and another part that flows in the dendrite, we have used several impedances (Fig. 1): *Z_a_*: the impedance of the extracellular medium in contact with the isopotential surface *S* (which touches the extracellular electrode and surrounds the intracellular electrode), *Z_s_*: the impedance of the current flowing through the soma membrane (in contact with this surface *S*), and *Z_d_*: the input impedance of the dendritic tree. When measuring the equivalent impedance, we have (*Z_a_* || *Z_s_* || *Z_d_*) ⊕ *R_g_*, where *R_g_* is the impedance between the ground and the first isopotential surface which comprises the neuron (Supplemental Information, Appendix A). Note that the isopotential surfaces are necessarily continuous, but may be more irregular in shape than that schematized.

The electric potentials *V_i_* and *V_e_* are taken at the inside and outside borders of the membrane, respectively, at the level of the soma and relative to a reference point outside the neuron. According to the law of generalized current conservation, we have 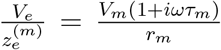. Note that the parameters *R_m_* = *r_m_* /Δ*S* and 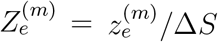 (where Δ*S* is an element of surface area of the membrane) are macroscopic parameters, while *r_m_* and 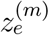 characterize the membrane at a microscopic level. This explains the difference between expressions 3a and 3b, because by definition we have *V_i_* = *V_m_* + *V_e_*. Note that if 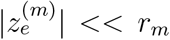, then expression 3b becomes equivalent to the input impedance of the dendrite (stick) in parallel with a portion of soma membrane (Fig. 1).

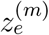 is the specific input impedance of the extracellular medium, as sensed by the membrane, as also defined by the generalized cable theory [22]. This parameter can also be applied to the intracellular medium, in case of a current source from an intracellular electrode, to take into account the impedance of the intracellular medium between the tip of the electrode and the membrane. In the open configuration, 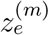 can be a resistance for a resistive extracellular medium, but can be more complex when taking into account effects such as polarization, ionic diffusion, capacitive effects, etc. Thus, the form of this frequency dependence contains the contribution from the extracellular medium. Note that the expression in Eq. 2 shows that, in the open configuration, the electrotonic length depends on frequency, even for a resistive extracellular medium.

Also, in a “closed configuration” (where the neuron is electrically a closed system, where all currents loop back to the neuron), for a resistive medium, we have 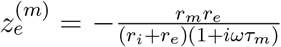 and 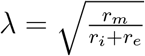. *Z_i_* = *r_i_* is the cytoplasm resistance per unit length and *r_e_* is the extracellular medium resistance per unit length, as defined in the Rall-Tuckwell model. In this simplified and resistive model of the neurons, one assumes that the extracellular current flows parallel to the axis of the dendrite. In this case, we see that λ does not depend on frequency. This is in accordance to the classic Rall-Tuckwell cable theory [23, 24]. In contrast, in an open configuration with resistive media (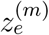 and *z_i_* are real positive), the modulus of λincreases with frequency (Eq. 2). Thus, in general, the electrotonic length (|λ|) of a ball-and-stick neuron depends on frequency, which is not the case in the Rall-Tuckwell model, as shown above.

An interesting consequence of this is that, if one measures the impedance 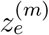 in a resistive configuration (for example in an isolated neuron embedded in ACSF), one should see the frequency dependence, which would validate the open configuration, as we will (see Results).

## 3 Results

We first describe the experimental results and the different experimental configurations, then we propose different models to fit the experimental measurements.

### 3.1 Experimental measurements

The experimental protocol consisted of *in vitro* whole-cell patch-clamp recordings of striatal neurons, either from primary cell cultures or in acute brain slices, while simultaneously recording the potential in the vicinity (5-10 *μ*m) of the neuron using a second reference electrode (Fig. 1A and 2B1). The set-up thus consists of three electrodes: the intracellular electrode, the reference electrode and the ground. Frozen white Gaussian noise is injected in the cell via the patch-clamp electrode, and is measured according to two configurations: either one measures the intracellular-to-cxtraccllular (intracellular electrode with respect to reference electrode) potential, or the cxtraccllular-to-ground (reference electrode with respect to ground).

**Figure 2:**
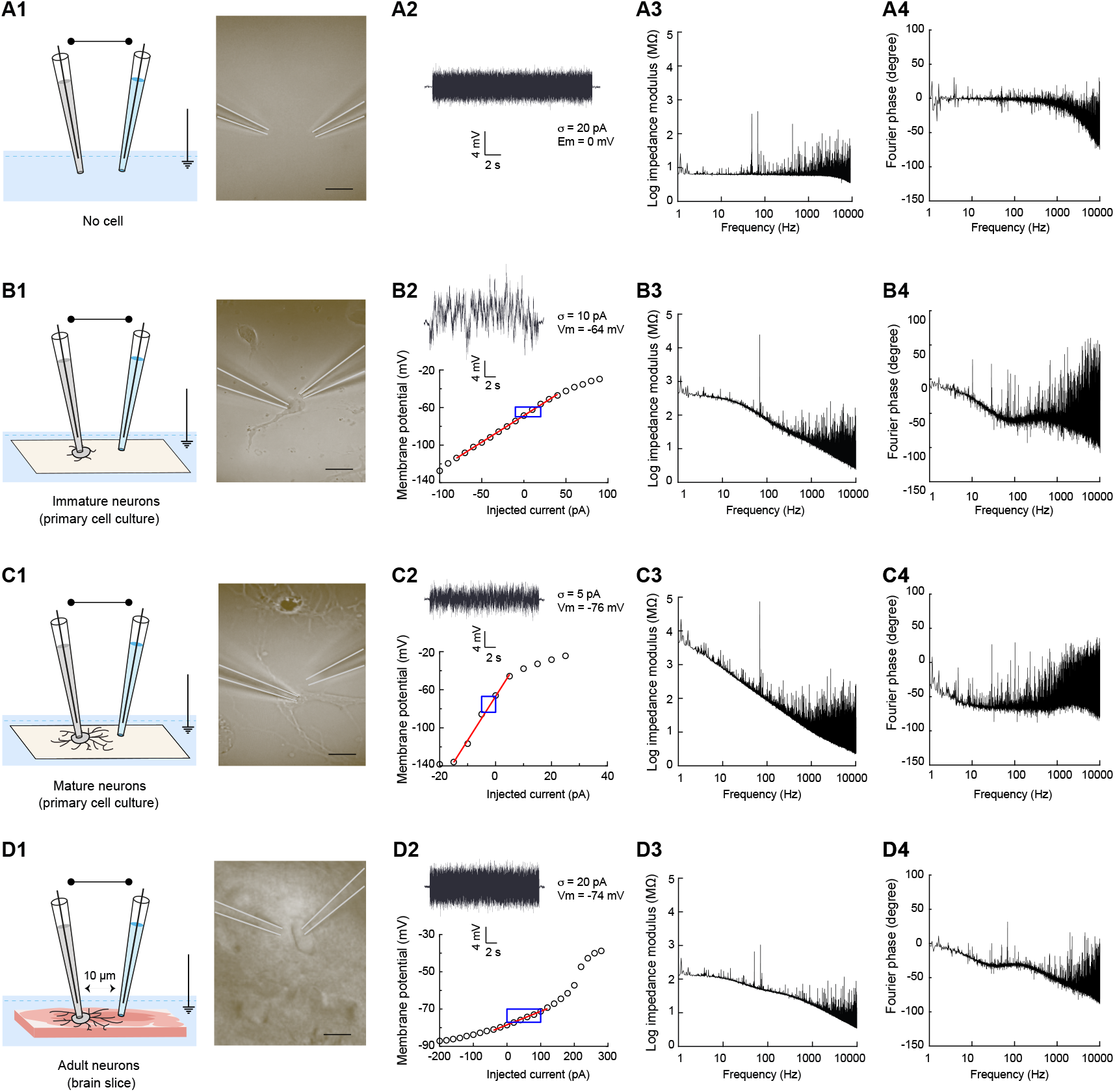
Impedance measurements in different preparations using a three electrode setup. **(A)** Measurement in ACSF. Description of the *(1)* experimental preparation (no biological sample is present in the ACSF solution) with *(2)* an example voltage response and for neuronal recordings, the corresponding I-V curve with the linear fit in red and the window of the voltage amplitudes during the current injection in blue. (3) and (4) show respectively the Fourier modulus and phase spectra of the voltage difference between whole-cell patch electrode and reference electrode as a function of frequency. **B,C,D**: same set-up for different experimental preparations: recordings were made in primary cell cultures of striatal tissue (day 3 to day 10 after seeding) from neurons with almost no neurites **(B)**, with extended dendrites **(C)** and from striatal neurons in acute horizontal brain slices **(D)**.

Several templates (from 1 to 71) of the same Gaussian white noise with a flat spectrum between 0 and 10000 Hz, were injected into each recorded neuron. Importantly, for the subthreshold range of voltage responses considered here (−67 mV ± 2.3 mV), the membrane I-V curve of striatal neurons was linear and only subthreshold responses to current injections (± 5 to 20 pA, adjusted according to the membrane resistance) were considered for analysis. More precisely, recordings, using the very same electrophysiological set-up, were obtained in four different experimental preparations (Figs. 2 and 3). We describe below the experimental results from the simplest experimental preparation in which two electrodes were added to a homogeneous ACSF solution without biological sample (Fig. 2A1), to a more complex preparation, in which recordings were obtained from neurons within an acute brain slice (Fig. 2D1). We also tested the influence of the dendritic arborization by recording either in non-arborized neurons with almost no neurites (Fig. 2B1) or in arborized neurons with extended dendrites (Fig. 2C1) in primary cell cultures. For each experiment, the Fourier modulus and phase of the voltage difference between the whole-cell recording electrode and the reference electrode were calculated and then averaged (as illustrated in Fig. 1). The same analysis was also applied to the voltage difference between the reference and ground electrodes (Fig. 3).

**Figure 3:**
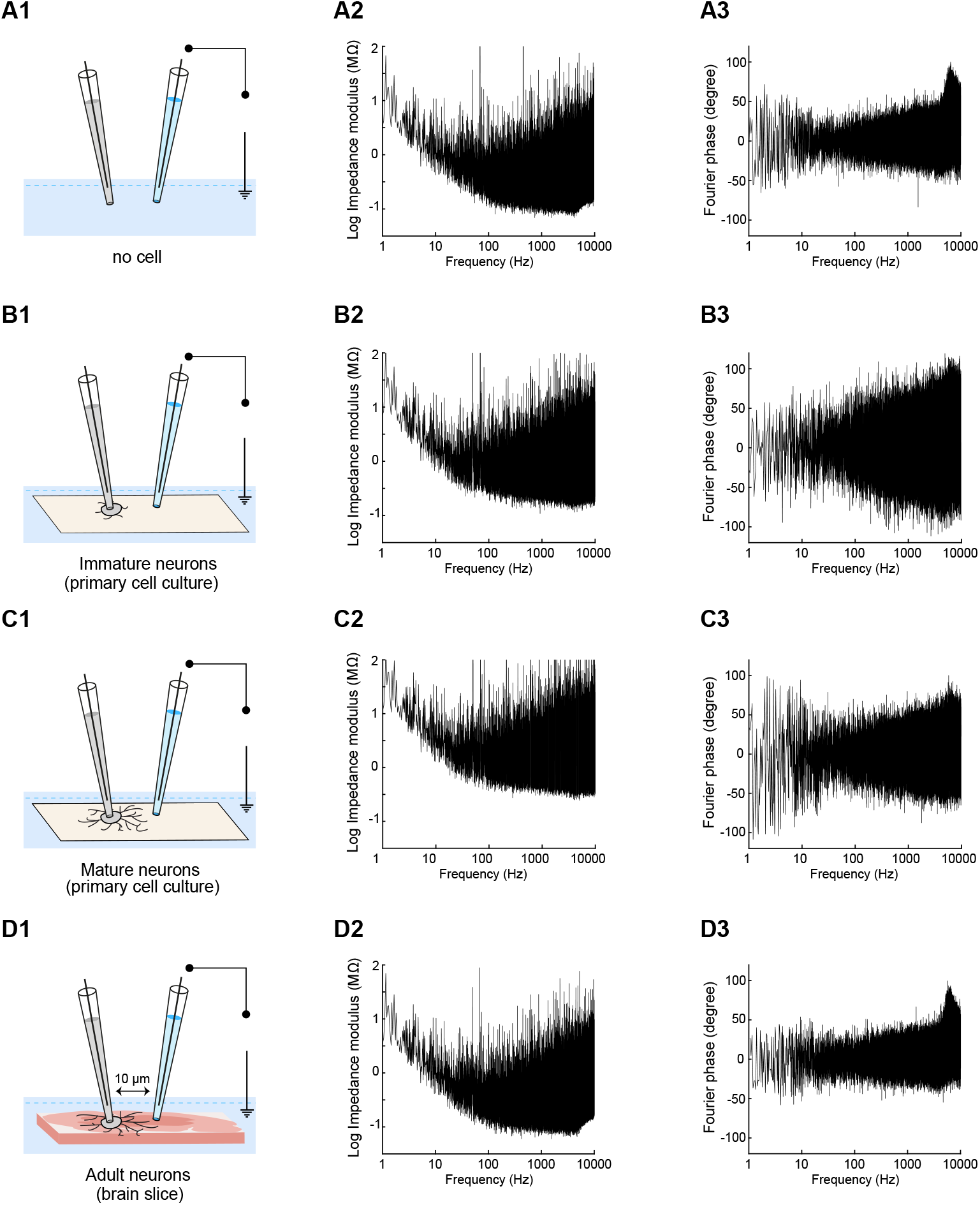
Measurements between the extracellular electrode and the ground (extracellular-to-ground impedance). Same arrangement of panels as in Fig. 2. Left: scheme of the recordings in the different preparations; Middle: modulus of the impedance; Right: phase of the impedance.

We describe below the experimental results from the simplest configuration of the two electrodes in ACSF, then neurons in a quasi-homogeneous medium^4^ (primary cell cultures), and finally a more complex configuration of neurons in acute brain slices.

### 3.2 Measurements in ACSF

In this section, we examine the simplest configuration consisting of two electrodes in a homogeneous medium solely constituted by ACSF (Fig. 2A and 3A). The glass pipettes containing the electrodes were situated at a distance of 5 to 10 *μ*m. Figure 4 shows the measured impedance between the two electrodes. The observed frequency dependence appears above 2 *kHz* for the modulus (Fig. 4A) and above 500 Hz for the phase (Fig. 4B). This frequency dependence of the impedance modulus is negligible for frequencies lower than 2 kHz, but not for higher frequencies because Δ*log*_10_*V* ≈ 50 %between 2 kHz and 10 kHz, similar to previous measurements using patch electrodes in the extracellular medium [6]. The phase of the impedance also shows a frequency dependence. It is negligible for frequencies smaller than 0.5 kHz but for frequencies between 0.5 and 10 kHz, it varies of about 50 degrees. Our interpretation is that these frequency dependences come from the capacitive effect between the two electrodes (which was estimated of *C* ≈ 3.25 *pF*). We found that this capacitive effect is present in all experiments shown in the next sections, for intracellular and extracellular recordings.

**Figure 4:**
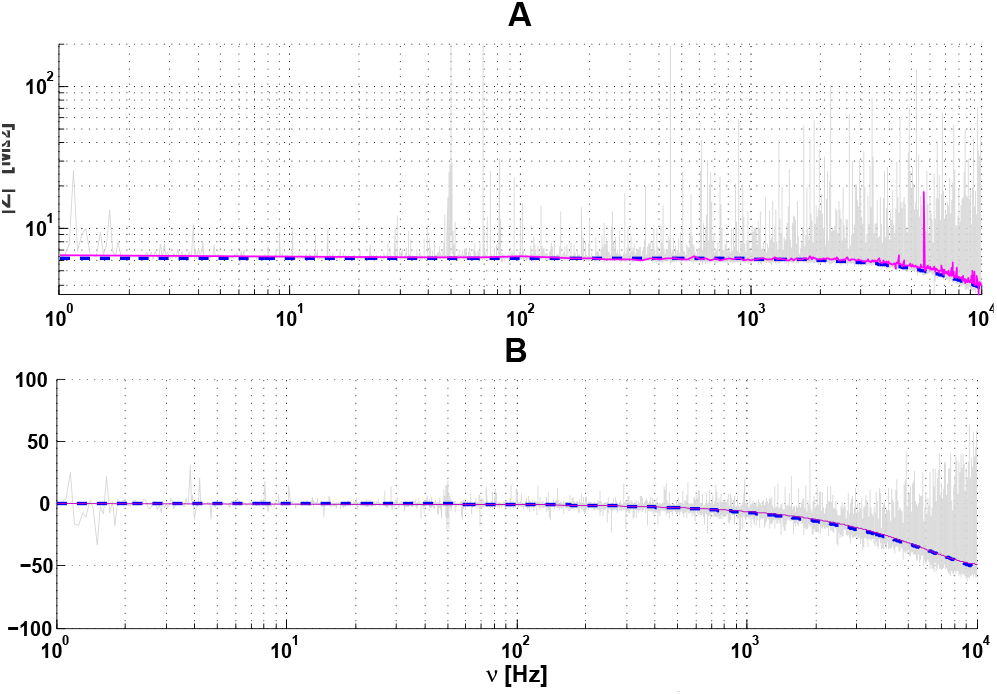
Impedance between two electrodes in ACSF. The distance between the two electrodes was of the order of 5 to 10 *μ*m, The modulus (A) and the phase (B) are represented as a function of frequency *ν*. A frequency dependence can be seen for *ν* > 2 *kHz* (A) and for *ν* > 500 *Hz* (B). The blue curve shows a capacitive effect of 3.25 *pF*. The magenta curve is a cubic spline fit of the experimental data (grey).

Importantly, in this part of the experiments, we observe a linear phase lag Φ = –*kν* on the phase Φ of the impedance, as if electrode polarization had a non-negligible impact on the measurement. However, we observe that the constant *k* is inversely proportional to the sampling frequency, which rules out a polarization effect (because with polarization, we would have Φ = –(*k* + *k_p_*)*ν* with *k_p_* independent of sampling frequency). Thus, the phenomenon of electrode polarization seems negligible here, contrary to Miceli et al. [6]. This difference probably occurs because we applied smaller amplitude currents (about 20 pA here, compared to 175-500 pA in [6]. Note that the linear phase lag was removed in Fig. 3A3.

According to Miceli et al. [6] and Wagner et al. [8], the electric conductivity of ACSF (“*ACSF_c_*” in Miceli et al.) is similar to the conductivity measured between two points in the extracellular medium (around 0.5 S/m). We did not evaluated the value of this conductivity here, because the very short distance between electrodes (5-10 *μ*m) induces capacitive effects. In this case, one would have to solve Laplace equation to estimate the conductivity from the macroscopic impedance measurement. In previous experiments such as [6], the inter-electrode distance (100-125 *μ*m) was sufficiently large to avoid capacitive effects.

### 3.3 Measurements in primary cell culture

#### 3.3.1 Non-arborized neurons

In this section, we first examine a relatively simplified system of immature neurons with little dendritic arborization, and laying in a simplified extracellular medium (primary cell culture), n=6 neurons (with few or no dendrites) were recorded in this quasi-homogeneous medium (Fig. 2B)

Figures 5, 6 and 7 show the fitting of different models to experimental measurements obtained with isolated cells immersed in saline, which is a medium that can be considered as homogeneous and resistive as a first approximation. The fitting to these measurements shows that the intracellular-to-extracellular and extracellular-to-ground macroscopic impedance clearly depends on frequency.

**Figure 5:**
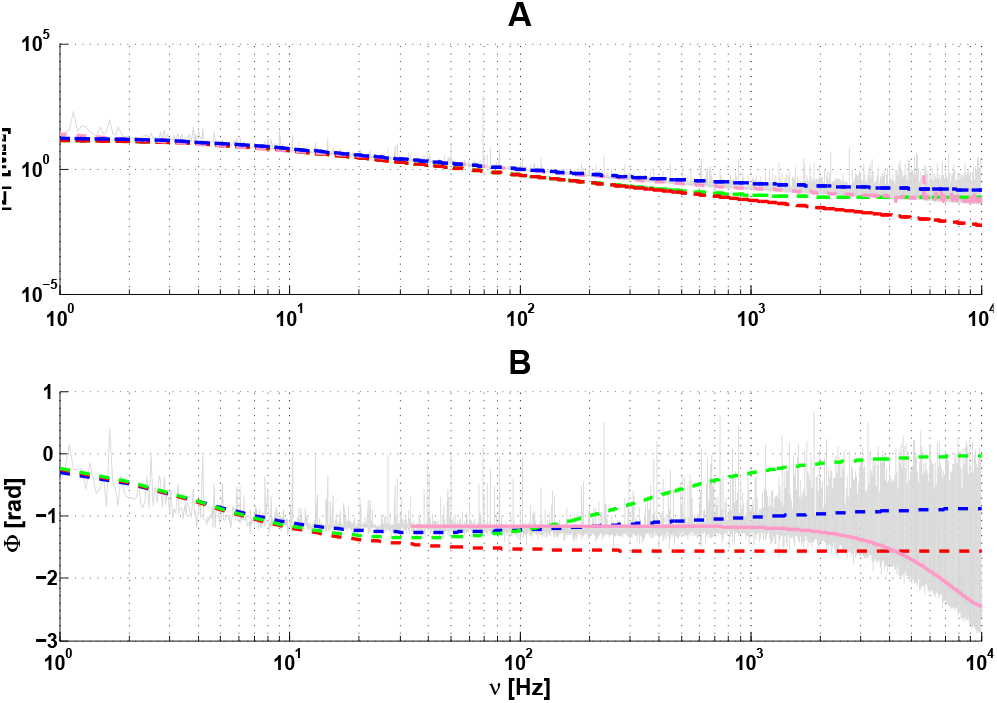
Cell-to-extracellular impedance *Z* as a function of frequency *ν* for a non-arborized neuron (no dendrites) in a primary culture (quasi-homogeneous medium). **A**: modulus of *Z* as a function of *ν*. **B**: phase of *Z* as a function of *ν*. The different models tested were, **Red**: (*R_m_*||*C_m_*) model, **Blue**: (*R_m_*||*C_m_*) ⊕ *Z_w_* ⊕ *R_asymp_*, **Gray**: experimental measurement, **Green**: (*R_m_*||*C_m_*) ⊕ *R_extra_* (Supplemental Information, Appendix B). The symbol ⊕ means “in series with” and the symbol || means “in parallel with”. Magenta: cubic spline fit of the experimental data. **Parameters**: *R_m_* = 810 *M*Ω, *τ_m_* = 30 *ms*; *A_w_* = 495 *M*Ω, *ν_wT_* = 0.1 *Hz*, *R_extra_* = 4 *M*Ω and *R_asymp_* ≈ 0.5 *M*Ω.

**Figure 6:**
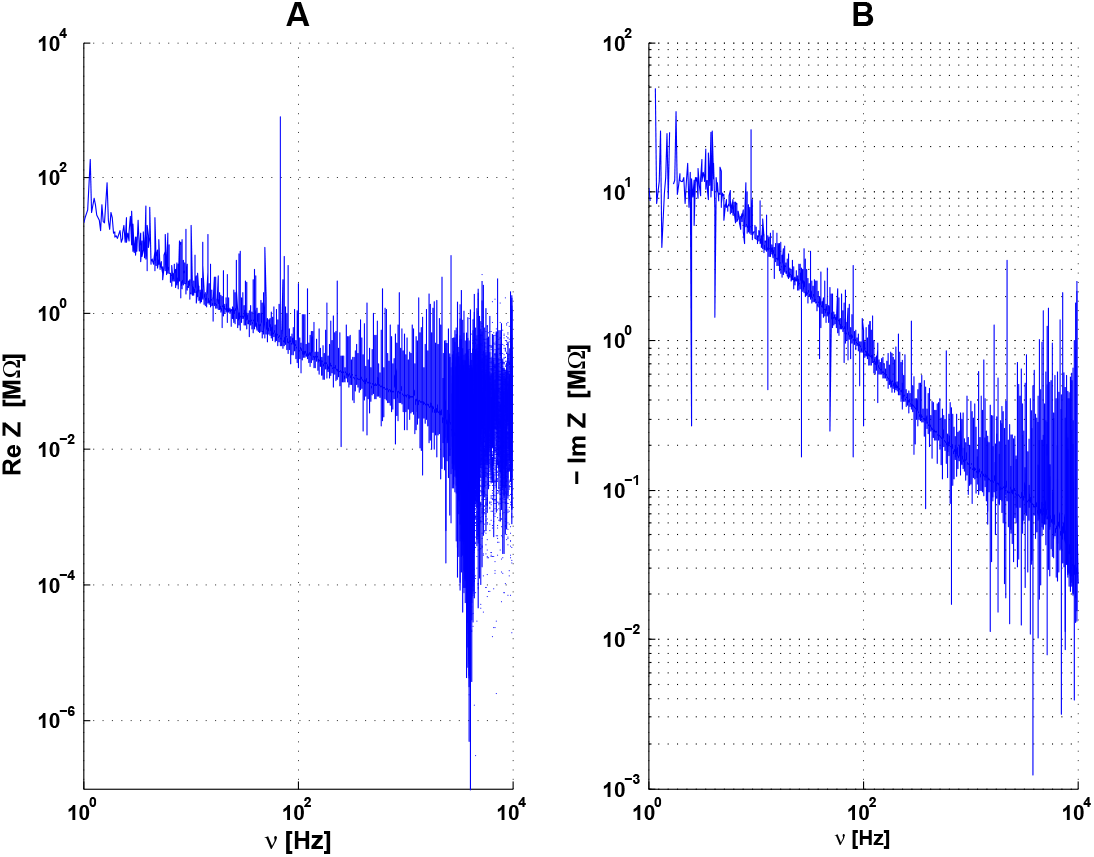
Impedance in polar coordinates for ccll-to-cxtraccllular measurements using experimental data from Figure 5 of the manuscript. A. Re(Z) vs. frequency from the data of A. B.-Im(Z) vs. frequency from the same data.

**Figure 7:**
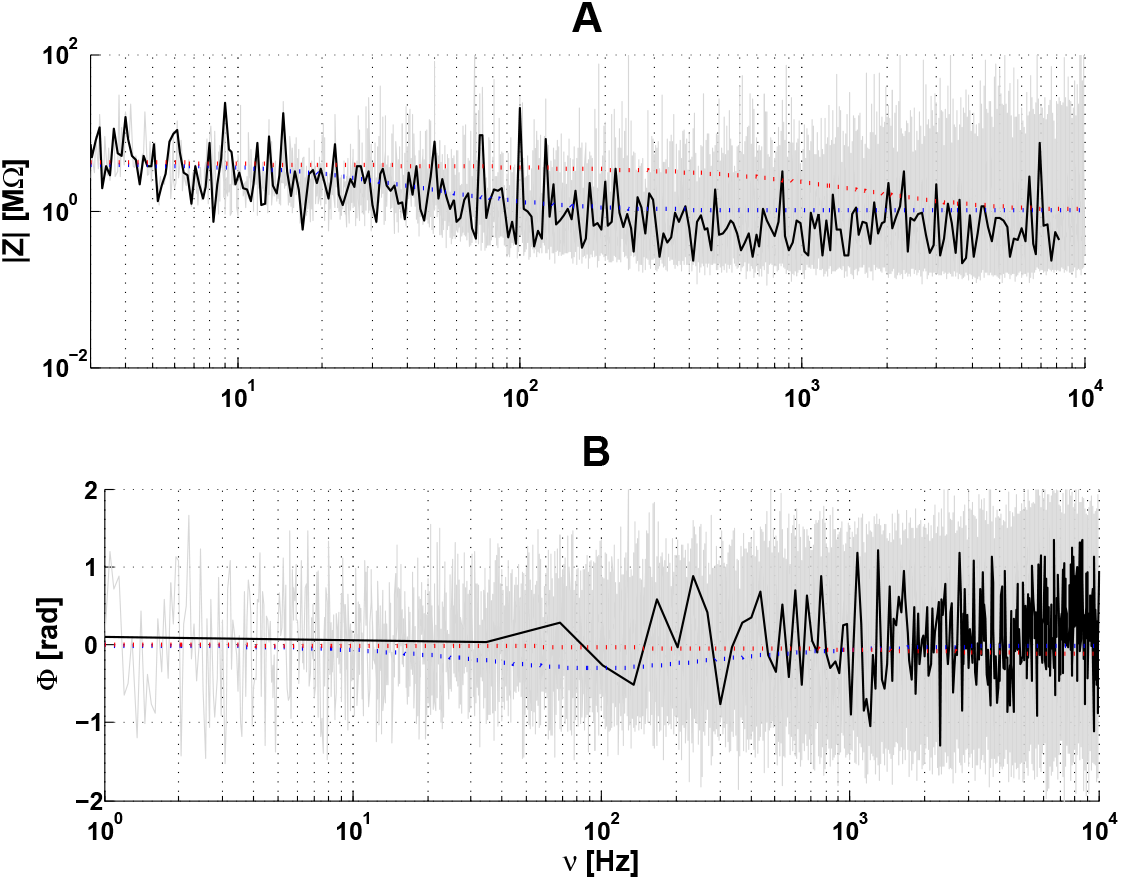
Extraccllular-to-ground impedance for a non-arborized neuron in culture. **A**: modulus of the impedance as a function of frequency. **B**: phase of the impedance as a function of frequency. Dotted lines correspond to the mathematical model given by Eq. 3b (Fig. 5 and Supplemental Information, Appendix A). The different fits shown are: Blue: **NMD**-Membrane + diffusive intracellular and extracellular media; Red: **NMR**-Neuron in a non-negligible resistive medium 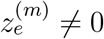; **Gray**: experimental measurement; **Black**: For **A**, cubic spline of the logarithm of the modulus and frequency of experimental data; For **B**: cubic spline of the phase and logarithm of frequency. **Parameters**: in all models, *R_m_* = 10 *M*Ω *τ_m_* = 25 *ms*, *r_d_* = 2 *μm*, *l_d_* = 100 *μm* and *Z_g_* = .1 *M*Ω; **NMR**: 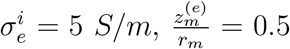, **NMD**: *ν_wT_* = 0.01 *H_z_*, 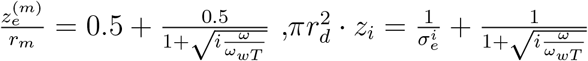.

Figure 6 also depicts the real and imaginary part of the impedance. Note that the instrumental noise on the phase is large compared to that on the modulus, the real |*Z*|*cos*Φ and imaginary |*Z*|*sin*Φcomponents are both more noisy than the modulus.

However, the representations of *Im*(*Z*)/*ν*(*Z*) and *Re*(*Z*)/*ν*(*Z*) complement well the analyses given in the paper.

The gray curve in Fig. 5 shows the measured impedance between the intracellular and extracellular electrodes. A RC circuit (membrane) in series with a resistance *R_medium_* (medium), modeling Region 1-2, cannot account for these impedance measurements (green curves in Fig. 5). Simulating the impedance with different values of *R_medium_* (Supplemental Information, Appendix B), could not mimic the experimental results. The red curve corresponds to *R_medium_* = 0, which is equivalent to consider that the membrane impedance is much larger than that of the extracellular medium. In this case also, it was not possible to properly fit the measurements.

The magenta curves in Fig. 5 are cubic spline fits of the experimental data. We observed a capacitive effect between the two intracellular and extracellular electrodes, as observed in ACSF, but larger. This indicates that the mean electric permittivity of the intracellular medium in Region 1-2 is larger than that of ACSF. This capacitive effect is responsible for a steep decrease of the phase values at high frequencies. This is consistent with a diffusive effect, because a resistive model with membrane would have a phase around *π*/2 *rad*. for *ν* > 100 *Hz*, which is difficult to reconcile with this phase measurement.

The gray curve in Fig. 7 shows the measured impedance between the extracellular electrode and the ground. A model with a simple dendrite (stick) with resistive intracellular and extracellular media could not account for these experimental measurements. The red curve corresponds to an extracellular medium with negligible resistivity compared to that of the membrane 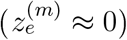. This case is equivalent to applying expression 3a. The green curve is such that 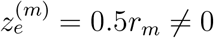.

In contrast, for the intracellular-extracellular measurement, we observed that the NMD model (*F*(*ω*) = 0 and 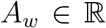; see Eq. 1) simulates (with some error) these experimental measurements with a macroscopic RC circuit in series with a resistance and a diffusive impedance (blue curve in Fig. 5). However, the residual error between experimental measurements and the diffusive model is very similar to what we observed for two electrodes in the ACSF (Fig. 4). This model also fits the extracellular-to-ground measurement, despite the noise on the phase, for a stick where both intracellular and extracellular Debye layers are modeled with a diffusive model (blue curve in Fig. 7). We propose an interpretation of these results in Section 3.3.3).

#### 3.3.2 Arborized neurons

We now follow the same approach as in the previous section, but in the case of recordings made from more mature neurons with extended dendritic arborizations, in primary cell cultures (n=6 cells), which corresponds Fig. 2C. Using the same scheme as in Fig. 5, we also investigate how the presence of dendrites influences the measured macroscopic impedance in this quasi-homogeneous medium.

By comparing the intracellular-to-extracellular impedances in Figs. 5 and 8, one can observe that the presence of dendrites has a negligible influence on the measured intracellular-to-extracellular impedance. The quality of the fit of the extracellular-to-ground impedance in Fig. 8 is very similar to that of Fig. 5. We considered a stick radius of 2 *μm* (as in Section 3.3.1) and a stick length of 600 *μm* (instead of 100 *μm*) to take into account the presence of a more extended dendritic arborescence (Supplemental Information, Appendix E). Note that the values of radius and length of the stick are not unique because the simulations can fit the experimental results equally well with a large number of dendritic parameters. However, the area of the dendritic stick is fixed when we fix the ratio 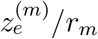, *r_i_* and *τ_m_* (see Eqs 3 and Eq. 2; for more details, see [22]). Thus, keeping the same stick diameter as in previous section, the fact that the length of the stick is much larger than in immature neurons, indicates that, in these experiments, the presence of dendrites has a much larger impact on the extracellular-to-ground impedance compared to the previous section. We have used a dendrite length which is 6 times larger for arborized neurons, which corresponds to visual inspection, but the results were weakly dependent on the exact value of this parameter (not shown).

**Figure 8:**
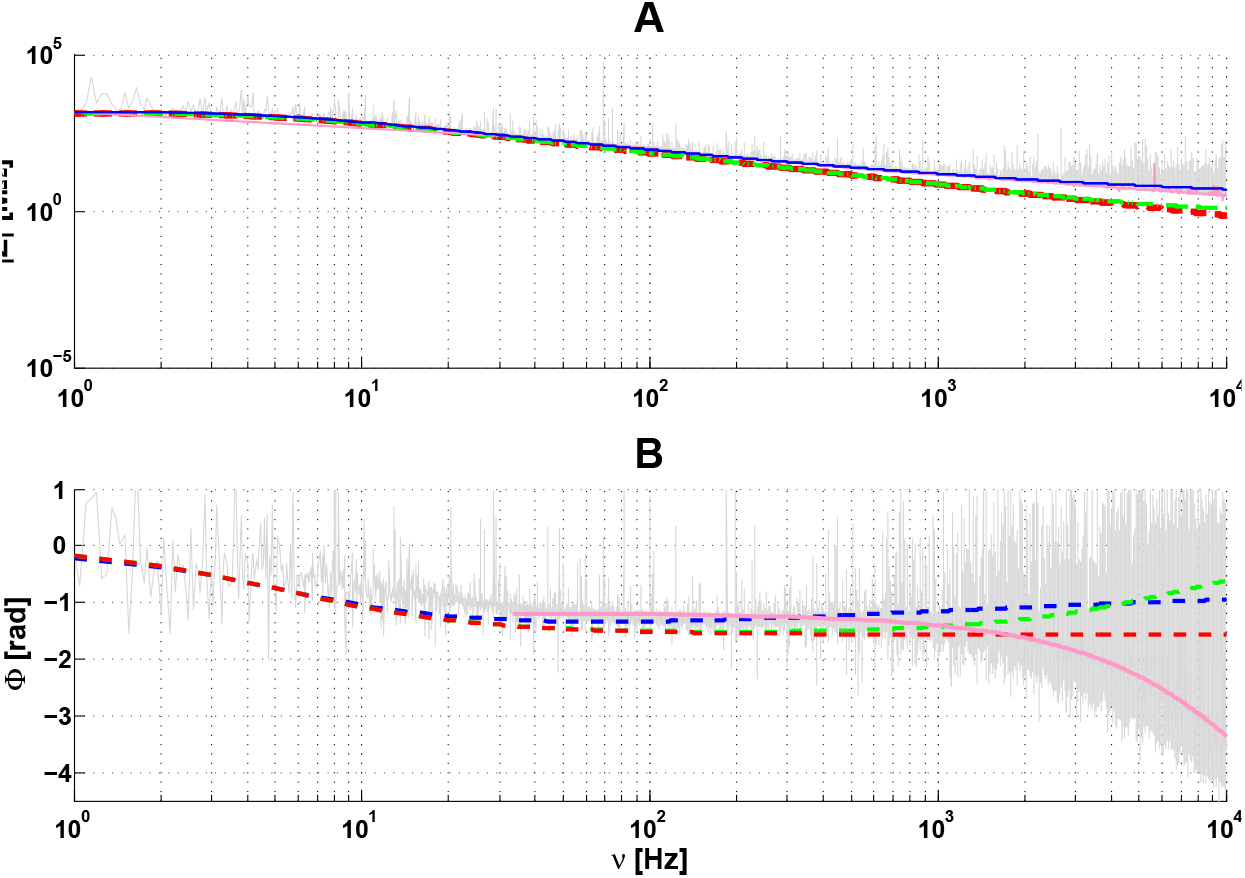
Intracellular-to-extracellular impedance *Z* as a function of frequency *ν* for an arborized neuron in a primary cell culture (homogeneous medium). Magenta: cubic spline fit of the experimental data. Similar description as in Fig. 5 with parameters *R_m_* = 1350 *M*Ω, *τ_m_* = 28 *ms*; *A_w_* = 450 *M*Ω, *τ_mw_* ≈ 0 *s*, *ν_W_* = 0.3 *Hz*, *R_extra_* = 1 *M*Ω, *R_asymp_* = 1 *M*Ω.

Thus, the fits of different theoretical models to experimental measurements of (arborized) neurons with dendrites give very similar results to those obtained in neurons without dendrites (non-arborized) examined in the previous section. In other words, the mean square error of the fit of the diffusive model is about 50 times smaller than that of the resistive model (with or without dendrites; Supplemental Information, Appendix G). The same capacitive effect as in ACSF was observed here for both arborized and non-arborized neurons (Fig. 4 in Section 3.2) and Fig. 5 and Fig. 8 in Section 3.3). It is thus important to stress that the 3-point measurement used here allowed us to directly measure the physical effect of the presence of dendrites on experimental measurements.

#### 3.3.3 Analysis of the experiments

The measurements from neurons in primary cell culture can be analyzed according to the electrical configuration depicted in Fig. 10. The isopotential surfaces indicated are central to our analysis. *S_i_* is the isopotential surface which corresponds to the potential measured at point *i*, where *S*_1_ corresponds to the potential measured by the intracellular electrode, *S*_2_ corresponds to the potential measured by the extracellular electrode and *S*_3_ corresponds to the ground potential. For simplicity, we will call Region *i* — *j* the domain delimited by surfaces *S_i_* and *S_j_*.

**Figure 9:**
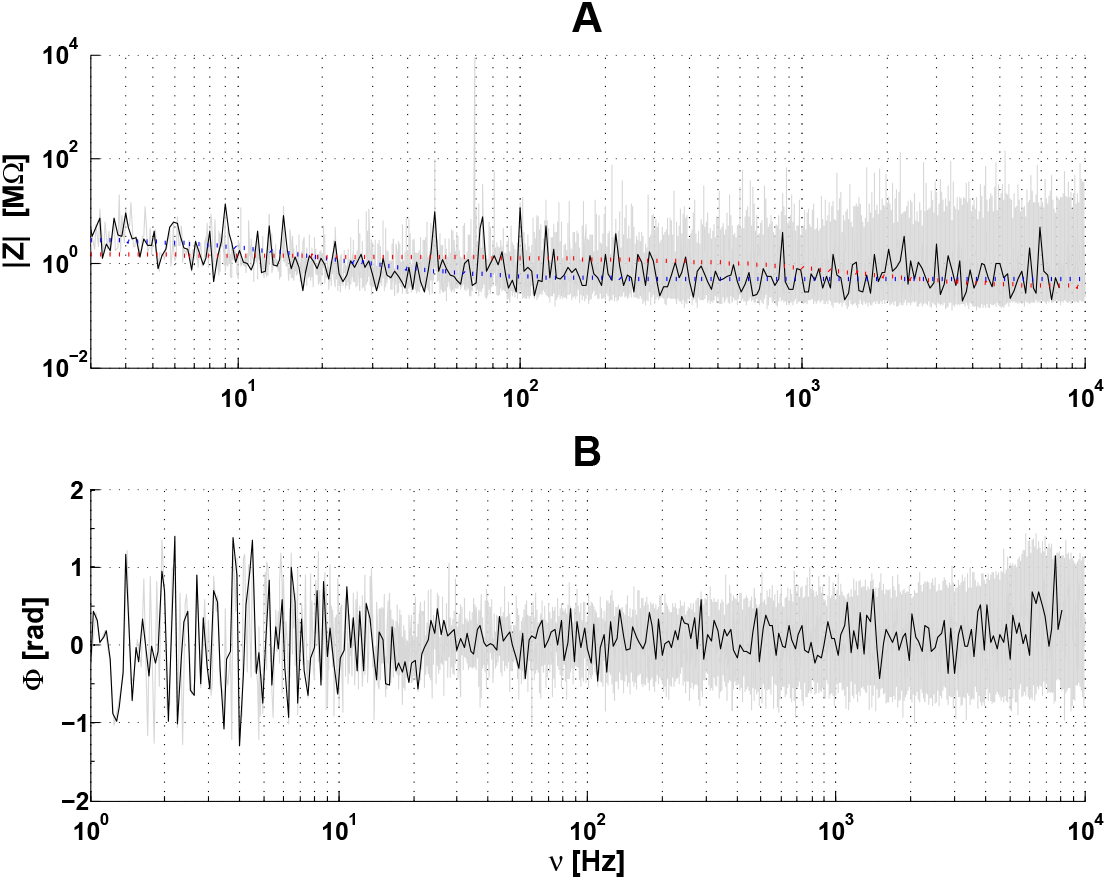
Extraccllular-to-ground impedance for an arborized neuron in culture. **A:** modulus of the impedance as a function of frequency. **B**: phase of the impedance as a function of frequency. Same color code as in Fig. 7. Dotted lines correspond to the mathematical model given by Eq. 3b (for details, see Fig. 7 and Supplemental Information, Appendix A). **Parameters**: *r_d_* = 2 *μm l_d_* = 600 *μm*, *R_m_* = 12 *M*Ω *τ_m_* = 35 *ms*, 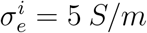 and *Z_g_* = 0.2 *M*Ω. **NMR**: 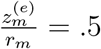; **NMD**: *ν_wT_* = 0.1 *Hz*, 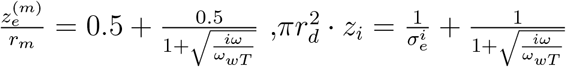.

**Figure 10:**
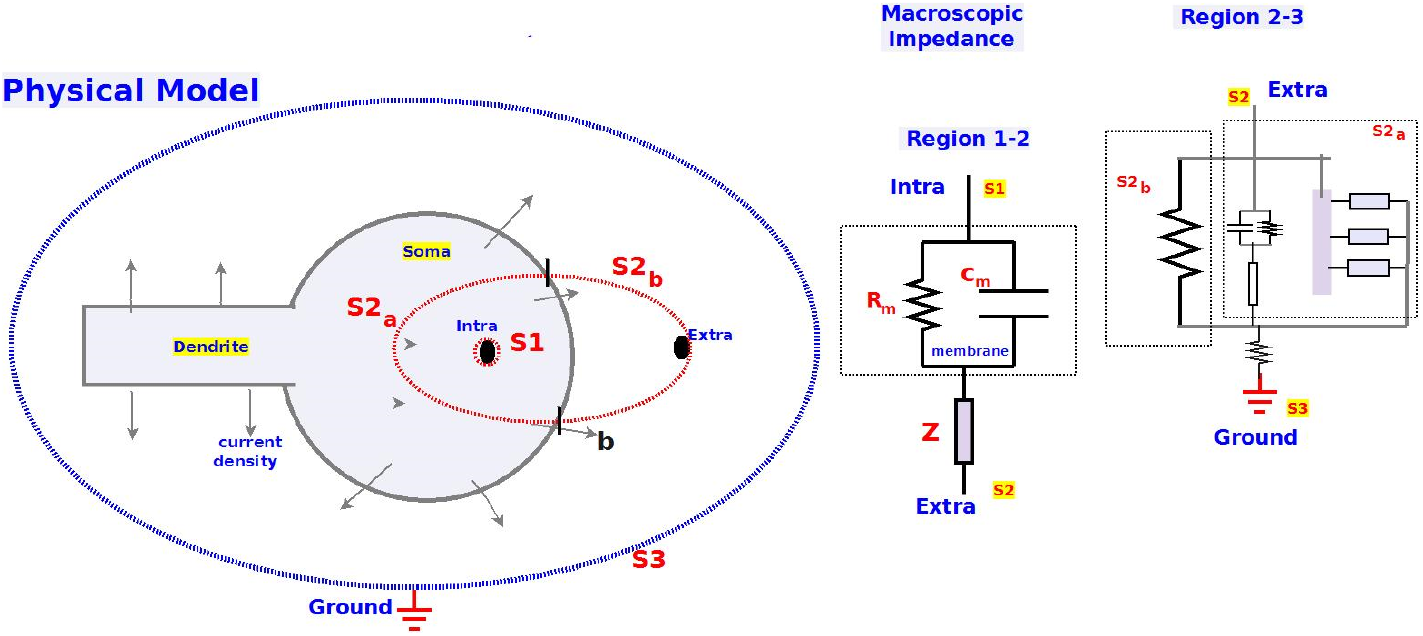
Electrical configuration of the cell and electrodes. Left: Physical model. The cell is schematized in gray, and the isopotential surfaces are labeled as *S*_1_, *S*_2_, *S*_3_. Right: equivalent electric circuits for the impedances calculated between the intracellular and extracellular electrodes, as well as between extracellular and ground electrodes. This corresponds to the isopotential surfaces *S*_1_ — *S*_2_ (Region 1-2) and *S*_2_ — *S*_3_ (Region 2-3), respectively. *S*_2_*b*__ is the part of surface *S*2 where the current directly goes to the extracellular medium, whereas *S*_2_*a*__ is the part of *S*_2_ where the current that passes through the intracellular medium before going extracellular (soma + dendrite). In other words, in Region 2-3, we have taken Eq. 3b where *Z_g_* is a resistance. *R_g_* is the resistance between the ground and the first isopotential surface which includes the neuron. Note that the drawing with equipotential surfaces (left) is a. schematic drawing where the only goal is to illustrate the link between equivalent electric circuits (right), the physical models and the experimental measurements (Supplemental Information, Appendix A).

First, because the extracellular medium surrounding the cell is mostly homogeneous and resistive, its macroscopic impedance cannot depend on frequency [10]^5^. We thus conclude that the surface *S*_2_ does not totally include the neuron, but cuts part of the soma, such that we have a portion of membrane over Region 2-3. Indeed, if the neuron was completely included into Region 1-2, then Region 2-3 would only consist of extracellular fluid, and would not exhibit (or exhibit negligible) frequency dependence, which was not what was observed (Fig. 7). This is in agreement with the fact that the intracellular and extracellular electrodes are very close to each-other (between 5 and 10 *μ*m), compared to the size of the soma (estimated around 15 *μ*m).

Second, the graph of the modulus of the impedance as a function of frequency in Region 1-2 (Fig. 5) is very different from Region 2-3 (Fig. 7). We thus conclude that the physical model is similar to the one represented in Fig. 10. Because the two points 1 and 2 are very close to each-other, we can simulate the impedance of Region 1-2 by an RC circuit in series with a diffusive impedance. However, the equivalent circuit of Region 2-3 cannot be the same because the current of the intracellular electrode divides into three parts: 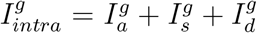, where 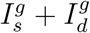 flows through the soma membrane and the dendrite, to the extracellular medium, while 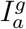 directly goes to the extracellular medium. Thus, the impedance of Region 2-3 is equivalent to expression 3b (Supplemental Information, Appendix A). Because this equivalent impedance is much smaller than the input impedance of the stick and of the soma membrane, the extracellular-to-ground impedance is also much smaller than the intracellular-to-extracellular impedance. Despite this small value, the frequency dependence of the modulus of this impedance can be clearly seen (Fig. 7). The experimental measurement of the impedance of Region 1-2 shows an additional capacitive effect similar to that seen in the previous section between intracellular and extracellular electrodes. However, this capacitive effect is negligible for the impedance measured in Region 2-3, which is coherent with the fact that the distance between the extracellular electrode and the ground is much larger than the distance between intracellular and extracellular electrodes.

Third, the growth of the modulus of the impedance of Region 2-3 for frequencies above 20 Hz is in full agreement with the diffusive model, as in Eq. 3b. This is not the case with the other models considered here. The minimum of the modulus is directly linked to the membrane time constant. The longer the time constant, the higher the frequency of the minimum. We estimate from the position of the minimum, a membrane time constant of about 30 ms, which is consistent with the membrane time constant estimated in this configuration. The growth of the modulus shows that 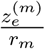 cannot be neglected (Fig. 5). The value of this ratio is estimated between 0.5 and 1 for null frequency (Supplemental Information, Appendix D).

Fourth, the values of the ratio *α* = *R_m_*/*A_w_* (model NmD for *ν* = 0) are approximately *α* ≈ 1 — 2 for both Region 1-2 and Region 2-3. Here, NmD stands for a model where each differential element of membrane is equivalent to a parallel RC circuit and where the intracellular and extracellular media are diffusive. These ratios do not correspond to a singular case, but were similar in all examined cells (Supplemental Information, Appendix G) in the same experimental conditions. Because neglecting *A_w_* would amount to have the NMR model of Fig. 5, these experimental results show that the intracellular and external Debye layers cannot be assimilated to a simple resistance. In contrast to NmD, the NmR configuration has resistive intracellular and intracellular media, but with the same membrane impedance. Moreover, the threshold frequency cannot be considered as infinitely large (*ν_wT_* = 0.1 *Hz* in Fig. 4 and *ν_wT_* = 0.001 *Hz* in Fig 5) because in this case, we would also have the NMR model. The threshold frequency *ν_wT_* of the diffusive impedance between surfaces *S*_1_ and *S*_2_ is greater than that between surfaces *S*_2_ and *S*_3_. This shows that surfaces *S*_1_ and *S*_2_ have very different curvatures. The small value of the Warburg threshold frequency (*ν_wT_* = *ω_wT_*/2*π*) indicates that the curvature radius of the isopotential surface *S*_2_ is much larger than the surface *S*_1_, which is consistent with the fact that surface *S*_1_ is at a zero distance from the tip of the intracellular electrode, which is not the case for surface *S*_2_ (Fig. 5; Supplemental Information, Appendix D).

Fifth, even for zero frequency, one does not observe the typical resistance of ACSF. If we calculate the resistance sensed by a fictive spherical source of diameter equivalent to the intracellular electrode (~ 1 *μm*), embedded in ACSF with an electric conductivity of *σ_e_* = 0.5 *S*/*m* [6], we have 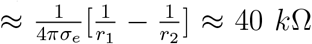. Here, *r*_1_ is the diameter of the Active source, and *r*_2_ is the average distance between the membrane and the spherical source. This value of 40 kΩ is smaller than the value in Region 1-2 predicted by the diffusive model for a near-zero frequency, because this value is much smaller than the estimated *A_w_*, which is about 30% to 45% of that of the membrane. This result is surprising, and different from the approximation usually made, assuming that the intracellular impedance is much smaller in amplitude compared to the membrane impedance for near-zero frequencies.

How to explain this result? According to the theory and experimental measurements of the electric conductivity of heterogeneous media [27, 28], the phenomenon of polarization occurs when an electric field is applied, and this contributes to lower the electric conductivity of the medium. Similar considerations apply to the phenomenon of ionic diffusion, which also lowers the conductivity [9, 22]. In the polarization effect, the electric field polarizes the intracellular medium, which determines an electric resistivity that is larger than the most conductive part of the medium. Thus, electric polarization diminishes the apparent electric conductivity in a heterogeneous medium, which directly affects the value of *A_w_* (Supplemental Information, Appendix C). Nevertheless, the present measurements show that the characteristic relaxation time of polarization (also called Maxwell-Wagner time), is negligible in these experimental conditions because the diffusive model alone can fit very well the experimental results^6^. If the polarization relaxation time was not negligible, there would be an additional frequency dependence to take into account in addition to the diffusive model. For example, if we consider diffusive model with polarization, and a non-negligible relaxation time, then we would have a supplementary frequency dependence relative to the diffusive model (not shown). Note that the diffusive model is equivalent to a model where the polarization relaxation time becomes very large when the frequency tends to zero (Supplemental Information, Appendix F). Because the diffusive model fits very well the measurements, adding polarization and or an additional resistance is not necessary. We conclude that the intracellular medium is such that the cutoff frequency *f_c_* = 1/2*πτ_mw_* of the low-pass filter due to electric polarization [11] is larger than 10 *kHz* in this experiment.

The large extracellular medium impedance can also be explained by the tortuous structure of the intracellular medium. Because the electric field lines cannot follow this tortuous structure, the charges that are moving in the extracellular space due to the electric field are subject to various obstacles [40]. To move away from these obstacles, ionic diffusion is necessary in addition to the electric field. Thus, it is expected that the impedance modulus of such a tortuous medium is larger than ACSF.

The high value of the impedance (high value of *A_w_*) is also due to the fact that the size of the soma is much larger than the plasmic membrane thickness. The ratio between the two is of the order of 1000, because the membrane has a thickness around 7.5 *nm* while the soma has a typical size around 10 *μm*. If we consider a slice inside the soma, we get approximately *R_soma internl_* < *A_w_*/1000 = 4 *M*Ω/1000 ≈ 4 *k*Ω which is much smaller than *R_m_*. We can repeat this for further slices in series of similar thickness. Because *A_w_* contains the contribution of external and internal Debye layers^7^, the value of *R_soma internal_* evaluated is smaller than 4 *k*Ω. Thus, the medium inside the soma is much more conductive than the membrane, as postulated by the standard model, but the difference is not as large as assumed by that model (red curve in Fig. 5).

Finally, we observed that the impedance measured between the extracellular electrode and the ground is much more noisy than the intracellular-to-extracellular impedance. This is, in part, due to the smaller amplitude of the latter impedance, and to the fact that the power falls off at higher frequencies, where instrumental noise becomes dominant. However, it should be noted that, besides the presence of this higher level of noise, the measurements also show that the modulus of the two theoretical diffusive impedances in Regions 1-2 and 2-3 are in full agreement with the experimental results. This indicates that the extracellular electrode is sufficiently far away from the membrane and does not disturb its ionic environment (Debye layers). It is nevertheless close enough so that the isopotential surface *S*_2_ only cuts a portion of the soma.

To conclude this analysis, the present measurements from neurons in primary cell culture cannot be made compatible with a resistive system, both for the modulus and phase of the impedance. However, both seem compatible with the impedance profile predicted by a diffusive system, which we interpret as being essentially due to the presence of Debye layers surrounding the membrane. However, to be rigorous, we must also mention the influence of the multiple obstacles inside and outside the neuron, which also have Debye layers and will influence the current flow. Importantly, the lack of major differences between the measurements from immature non-arborized neurons and those obtained from neurons with an extended dendritic tree argues against the hypothesis that the presence of dendrites could have a significant impact on impedance measurements.

### 3.4 Measurements in acute brain slices

In this section, we present and analyze impedance measurements of mature and fully-arborized neurons (n=9 cells) recorded in acute brain slices (Fig. 2D). In this case, the tissue surrounding the neuron is quasi-intact, and the extracellular medium is the neuropil, which is very heterogeneous [41]. We analyzed the effect of such an environment on the frequency dependence of the macroscopic impedance.

We used the same experimental method (3-point recording) as in previous sections. We observe a different frequency dependence for the impedance of Region 1-2 (compare Figs. 5, 8 and 11). This shows that the presence of a complex extracellular medium (neuropil) causes more complex effects in the vicinity of the neuronal membrane. However, if we consider two different threshold frequencies for intracellular and extracellular media, we obtain

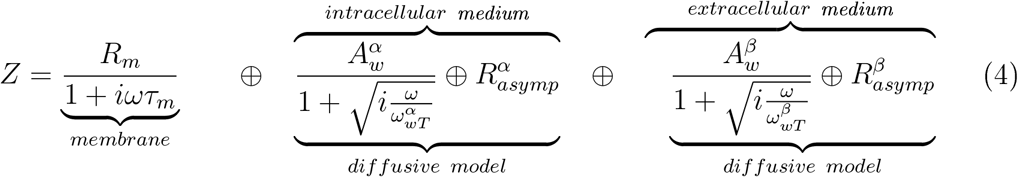

which gives excellent fits (Fig. 11). Note that we have kept the characteristics (parameters) of the previous experiments for modeling the intracellular medium.

**Figure 11:**
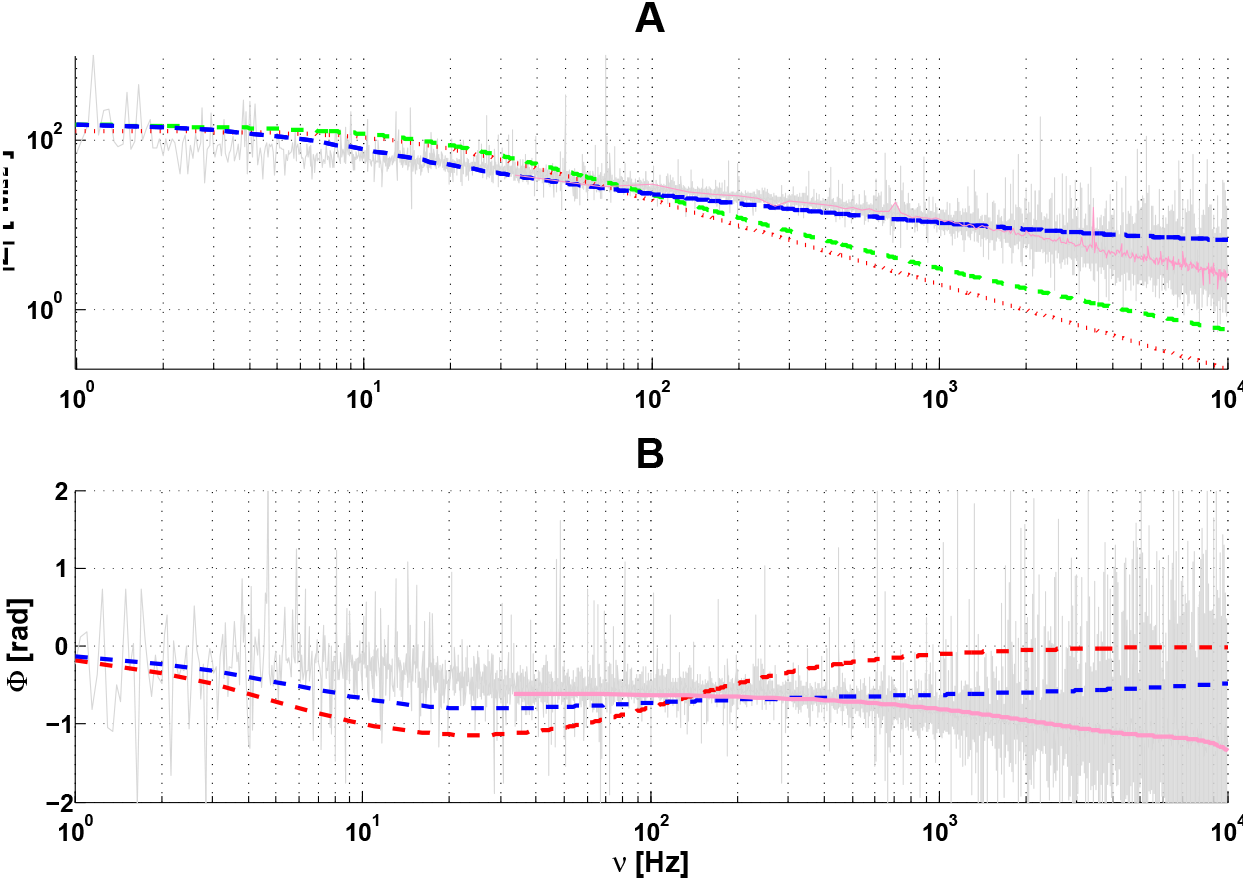
Intracellular-to-extracellular impedance *Z* as a function of frequency *ν* for a neuron recorded in acute brain slice. **A:** modulus of *Z* as a function of *ν*. **B**: phase of *Z* as a function of *ν*. **Parameters**: *τ_m_* = 10*ms*, *R_m_* = 128 *M*Ω; 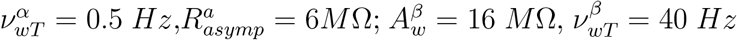 and 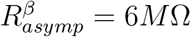. The color code is **red**: fit with resistive intracellular and extracellular media (16 *M*Ω); **green**: fit with a diffusive intracellular medium (a) and a resistive extracellular medium (16 *M*Ω); **blue**: fit diffusive intracellular and extracellular media (Eq. 4); Magenta: cubic spline fit of the experimental data. The latter case best fits the experimental measurements. The mean square error of the diffusive model is about 3-fold less that than of the resistive model (Supplemental Information, Appendix G).

In Region 2-3, we observed a similar frequency dependence as in previous configurations (compare Figs. 7, 9 and 12). In the case of the diffusive model, we used a model of the macroscopic impedance *Z_b_* (impedance of the extracellular medium as sensed by Surface *S*_2_; Fig. 10), which was taken from a previous study [9], with the difference that we added a series resistance (8 *M*Ω). This addition is justified in Supplemental Information, Appendix C (Eq. C.5). This impedance is necessarily macroscopic because it is given by the ratio of the measured current between two points separated by macroscopic distances (as defined by the region in between the two isopotential surfaces going through each point). Accordingly, we used 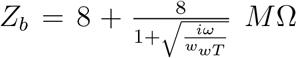 where *ν_wT_* = 40 *Hz*, as determined in [9]. Note that the particular choice of the parameters (*A_w_* = *R_asymp_* = 8 *M* and *w_wT_*) can be varied (by approximately ± 50 %) with no qualitative change in the results (Supplemental Information, Appendix G). While many combinations of parameters fit the data for diffusive models, we did not find a single parameter set of resistive models that could fit the data.

**Figure 12:**
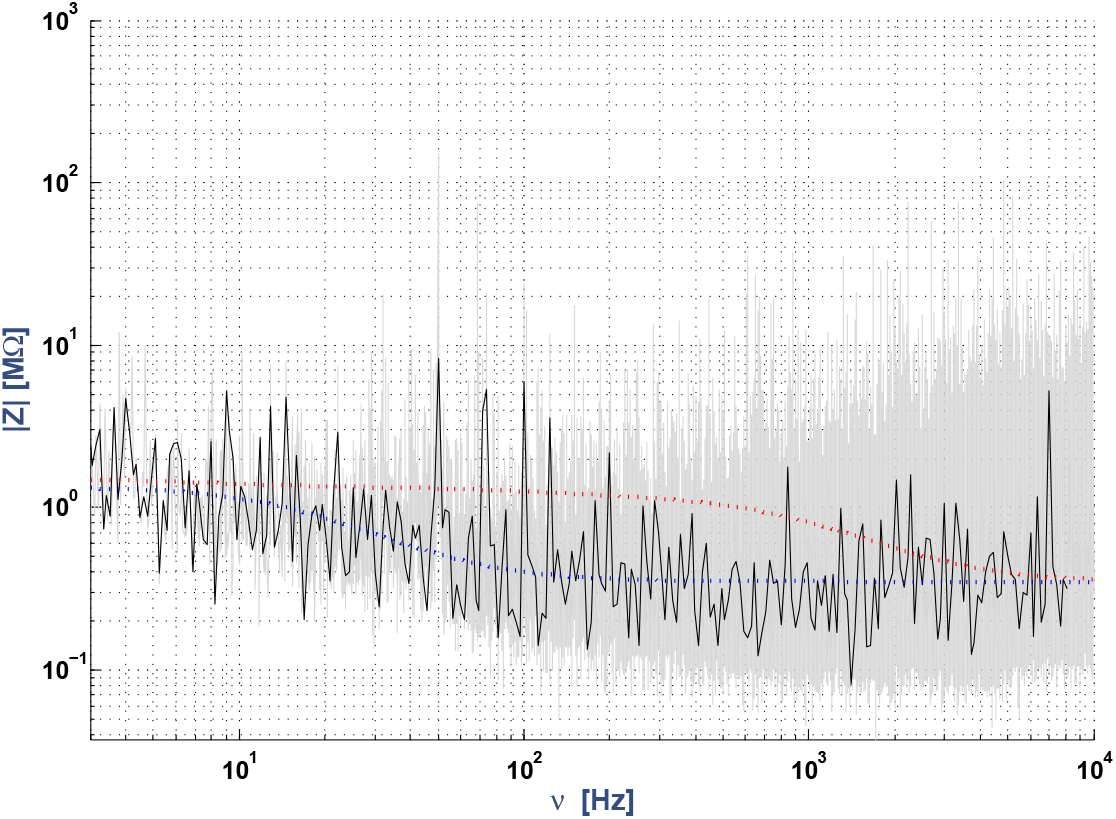
Extracellular-to-ground impedance for a neuron recorded in acute brain slice. The same color code as in Fig. 7 was used. Dotted lines correspond to the mathematical model given by Eq. 3b (for details, see Fig. 10 and Supplemental Information, Appendix A). The modulus of the impedance is represented as a function of frequency. **Parameters**: *R_m_* = 5 *M*Ω, *τ_m_* = 10*ms*, *r_d_* = 2 *μm*, *l_d_* = 600 *μm*, 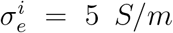 and *Z_g_* = .2 *M*Ω. **NMR**: 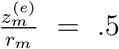; **NMD**: *ν_wT_* = 0.01 *Hz*, 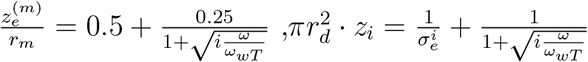.

The modulus of the extracellular impedance between the extracellular electrode and the ground (Region 2-3) in acute brain slices can be different from that in primary cell culture. The impedance sensed by the current source that does not flow through the dendrite and the cell membrane (Fig. 10) on Region 2-3 (between the extracellular electrode and the ground) are experimental conditions similar to extracellular measurements performed in other studies [5, 6]. Thus, for Region 2-3, we have chosen a similar model as for the experiments in primary cell culture.

#### 3.4.1 Analysis of the brain slice experiments

In the acute brain slice preparation, the observations are similar to the observations in primary cell culture, except that the best fit is obtained here when we use different threshold frequencies for intracellular and extracellular media. We now focus on the analysis and interpretation of such different threshold frequencies.

According to Gomes et al. [9], the extracellular impedance in brain slices has a threshold frequency between 40 and 60 Hz. Compared to the present experiments, the thickness of the extracellular medium was much larger in [9], because the extracellular electrode was located at a distance about 10 times larger. Indeed, in [9], the macroscopic impedance of the extracellular medium was larger than that of the intracellular medium. Thus, the present results, together with Gomes et al. [9], suggest that the threshold frequency of the extracellular medium is larger than that of the intracellular medium. This larger threshold may indicate that the medium is more tortuous inside the cell compared to the extracellular medium.

This result is also in qualitative agreement with the experimental measurements of Gabriel et al. [7] and Wagner et al. [8] because the apparent conductivity and permittivity of the diffusive model are respectively very low and very high, which gives a very large dielectric relaxation time (Supplemental Information, Appendix F). According to the experimental results of [7], the electric permittivity of the extracellular medium is much larger in the neuropil, compared to that of the extracellular fluid in cultures. The value of the permittivity is in agreement with the experimental measurements [7, 8] and is estimated to be 10^5^ to 10^7^ times larger than that of ACSF. Notice that the results of Wagner et al. were obtained *in vivo* and are not identical to that of Gabriel et al. obtained *in vitro*. This difference may be due to the fact that the linear approximation of the medium impedance may be valid *in vitro* but not *in vivo*. Can the linear approximation be considered as a first estimate? Is the deviation due to the presence of ongoing (spike) activity? These questions await experimental testing.

The correspondance between the theoretical models presented in Section 2.2 and the different experimental configurations is summarized in Table 1.

**Table 1:**
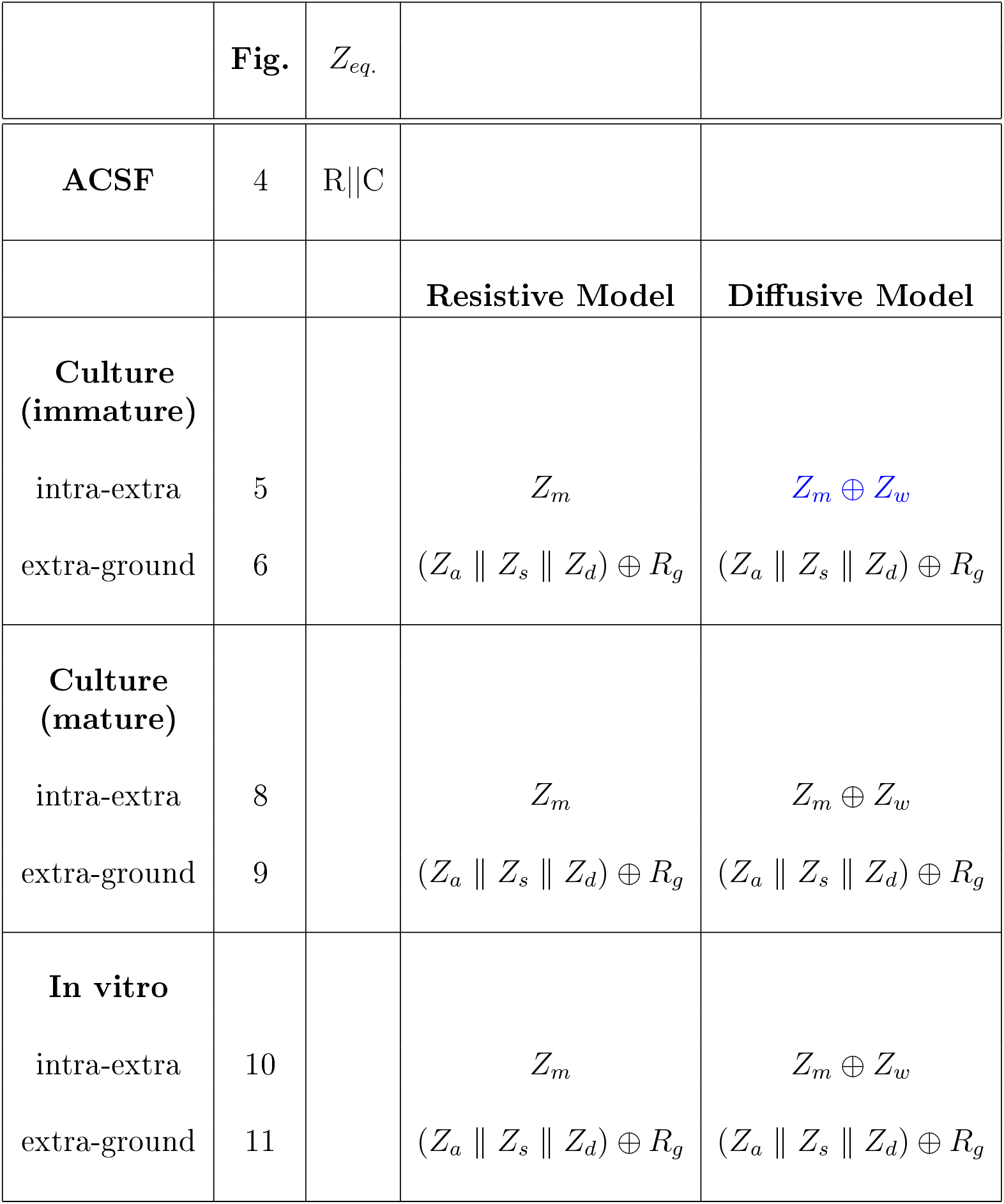
Table showing the different biophysical models used for fitting the experimental data in the different figures of the manuscript. **1:** *Z_m_* = *R* || *C* corresponds to the impedance of a portion of membrane where *R* and *C* are respectively the resistance and capacitance. **2:** *Z_w_* is given by expression 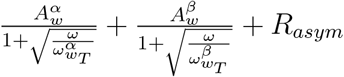 where 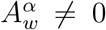 and 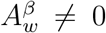 and *ω_wT_* ≠ ∞. In primary cell cultures, we have used 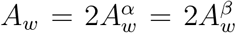 and 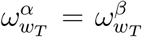. In the *in vitro* conditions, we have used 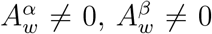 and 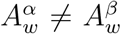 with 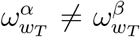 and different from infinity. *Z_a_* is the impedance of the region defined by the isopotential surface 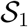 passing through the extracellular electrode and the first isopotential surface 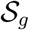 that totally includes the neuron, *Z_s_* is the impedance of the part of the soma included between 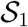 and 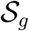, *Z_d_* is the input impedance of the dendrite bet ween surfaces 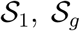, and *Z_g_* is the impedance of the region between 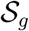 and the ground. *Z_g_* = (*Z_a_* || *Z_s_* || *Z_d_*) ⊕ *R_g_* where || means “in parallel” and ⊕ “in series” (Fig. 1).

## 4 Discussion

In this study, we provide for the first time, impedance measurements around neurons, in different experimental preparations and a comparison between extracellular and intracellular impedance measurements. The aim was to determine which is the most plausible physical model that accounts for the electric properties of the extracellular medium. We used an experimental measurement consisting of three points, an intracellular electrode, an extracellular electrode in the close vicinity of the membrane (a few microns), and a reference electrode (ground) located far away. Using the same recording set-up, we considered different preparations ranging from acute brain slices, primary cell cultures with neurons of different morphological complexity, and measurements in ACSF solution as control. In addition to these measurements, we have provided a detailed fitting of the measured impedance using several biophysical models of the extracellular medium. The main conclusions are that (1) in no circumstance, the resistive model is able to fit the whole data set; (2) the measurements point to a dominant role of ionic diffusion, as soon as a membrane is present; (3) the measurements suggest that the underlying mechanism is the ionic diffusion associated to the Debye layers around the membrane; (4) additional capacitive effects may be needed to explain the differences between experimental preparations.

Regarding the first conclusion, a resistive medium cannot account for any of the measurements, except when electrodes are placed in ACSF. The resistive nature of the impedance measured in ACSF is expected because this saline solution is the simplest case of a resistive medium. In this case, we had to consider an additional capacitive effect in parallel with the two electrodes (Section 3.2). This shows that the measurements with the micro-pipettes do not create an apparent frequency-dependence of the measured medium, and once the capacitive effect is removed, one recovers the correct resistive measurement in this case.

The second conclusion, that ionic diffusion plays a prominent role, is supported by our fitting analysis considering different model alternatives. This suspected presence of ionic diffusion (and the associated Warburg impedance) agrees with previous studies showing a role for ionic diffusion. Macroscopic measurements of the impedance of brain tissue [7, 8] showed a frequency dependence of the electric parameters which is consistent with ionic diffusion, as pointed out by a theoretical study [20]. This study developed a mean-field formalism of Maxwell equations, which was necessary to properly account for macroscopic measurements that imply averages over large spatial volumes. In this mean-field framework, the predicted frequency scaling of ionic diffusion was found to be consistent with the experimental observations. It was further shown, using a two-electrode measurement setup with intracellular and extracellular electrodes, that ionic diffusion also accounted for the observed frequency dependence [9]. However, as pointed out in [16], the dendrites were neglected in our previous study. We provide here new data on this issue by showing that there was little difference between the impedance measured in intact (arborized) neurons or in (non-arborized) neurons with greatly simplified dendritic morphology. Therefore, the presence of dendrites is not a plausible cause to explain the deviations from resistivity.

The third conclusion, that the underlying mechanism is the ionic diffusion in Debye layers around the membrane, is mostly supported by the experiments in primary cell cultures where no intact neuropil is present. In these conditions, the medium is almost devoid of glial cells or neighboring neurons, and can be considered close to homogeneous saline. We also observed a frequency dependence in such conditions, when an intracellular recording was present, but not in saline with two electrodes in ACSF (Section 3.2). Therefore, we find that the presence of a membrane induces a frequency dependence, which we attribute as mainly due to the presence of Debye layers around the membrane. Debye layers not only constitute the basis of the membrane capacitance, but they also are characterized by ionic diffusion which participates to maintain the membrane potential. Our interpretation is that, when current flows from the intracellular electrode, the corresponding ions induce local concentration changes in Debye layers, which will re-equilibrate by ionic diffusion. This increases the modulus of the impedance and introduces a frequency dependence which signature can be seen as a diffusive (Warburg) impedance. Importantly, this impedance is a linear approximation, and thus, cannot capture the ratio *V*/*I^g^* for large variations of the membrane potential (such as during spikes; see discussion in [20]).

The fourth conclusion is that it was necessary to include capacitive effects to account for differences between the different preparations. These effects were not necessary for a cell in a homogeneous medium (in culture), but were required to fit acute brain slice conditions when the the electrodes were very close (about 10 *μ*m). On the other hand, no additional capacitive effect was seen between the extracellular electrode and the ground. This suggests the possibility that both intracellular and extracellular media have a significant diffusive component. We could fit the measurements assuming a different cutoff frequency between the media (about 5 Hz intracellular, and 40 Hz extracellular). This difference suggests that the medium is more tortuous intracellularly compared to the extracellular medium.

Importantly, the present results seem in agreement with the principle of least constraint of Gauss, according to which the introduction of constraints modifies the least as possible the movement of a system. In electromagnetism, the application of this principle means that the majority of charges will follow the path with the lowest impedance^8^. In Logothetis et al. and Miceli et al. measurements [5, 6], the magnitude of the minimal extracellular impedance would be of the same order as ACSF, which is a similar situation as our measurement between the extracellular electrode and the ground. In Gomes et al. [9], as in the present paper, the modulus of the minimal impedance between the intracellular and extracellular electrodes would be much larger because the charges are constrained to flow across the cell membrane, and not just flowing exclusively in the extracellular medium.

Taking together our results in different preparations, we conclude that, for the frequency range of electrophysiological phenomena (frequencies *ν* < 10 *kHz*), there is a very significant frequency dependence of the electric parameters in intracellular and extracellular media. Our experiments show that this significant frequency dependence should be taken into account when current flows across a cell membrane. On the other hand, there is very little frequency dependence if the current does not flow across a membrane^9^. This major difference suggests that impedance measurements should necessarily give different results if they are performed extracellularly, or using an intracellular recording. This could potentially reconcile contradictory measurements and also answer some of the questioning about the origin of the LFP signal [42].

This raises the question of which impedance is the most physiologically pertinent, the impedance of the medium alone, measured with extracellular electrodes, or the impedance between intracellular and extracellular media ? If the point of interest is to relate neuronal activity (ionic currents) with the extracellular potentials, then the relevant impedance is the intracellular-extracellular impedance as measured here. Indeed, the ionic currents in the membrane have to flow through a complex environment (Debye layers, various obstacles), which are important to generate the extracellular potential (which “sense” the current *after* its interaction with the complex environment). The impedance measured here correctly captures this effect and is the closest to the natural conditions. On the other hand, impedances measured with extracellular electrodes inform about how the extracellular medium reacts to currents injected extracellularly and does not reflect the situation with natural (membrane) currents. Thus, our study complements previous extracellular measurements, and goes further in biophysical realism by measuring the impedance pertinent for the genesis of extracellular potentials.

Our results may call for a redefinition of the classic “neuronal dipole” mechanism underlying LFP or EEG signals. However, such an extrapolation is not immediate, because our measurements were made in the linear regime, while the LFP and EEG signals are generated by active networks which involve nonlinear phenomena such as action potentials. Nevertheless, our experiments indicate that LFP and EEG signals should be affected by the frequency filtering measured here, because the formation of the neuronal dipoles will depend on currents flowing through the membrane/Debye-layers complex. In other words, the currents forming the neuronal dipole should include the frequency filtering due to the flow through the membrane/Debye-layers complex, and which can only be revealed by intracellular measurements.

## Author contributions

CP and LV did the experiments, AD and CB conceptualized the study, CB did the analysis, and all authors contributed to interpret the analysis and to write the manuscript.

## Acknowledgements

We thank Peter Vanhoutte, Nicolas Heck and Lisa Huet for help with the cell cultures. Research supported by the CNRS, INSERM, Collège de France, the European Community (grants H2020-785907 and H2020-945539) and the ANR (grant PARADOX).

## Supplemental Information

## Appendices Physical meaning of the method used to measure the impedance

In our experimental situation, we inject a current which is time-dependent, in a linear medium. In such conditions, the potential (relative to ground) is given by the general relation:

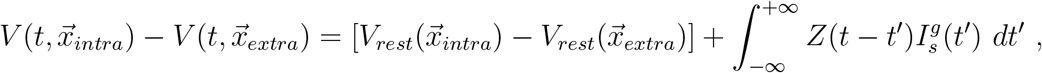

where 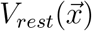 is the resting potential at position 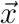 (at rest, 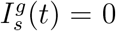). It follows that, in Fourier frequency space, the potential between intracellular and extracellular electrodes is given by:

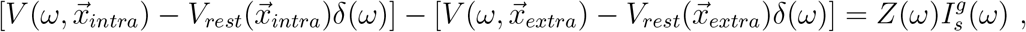

where *δ*(*ω*) is the Dirac distribution. Thus, we can write:

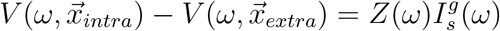

when *ω* ≠ 0. For *ω* ≠ 0 we have *δ*(*ω*) = 0 and thus 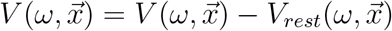 because 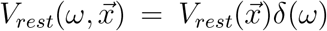. It follows that, if *ω* ≠ 0, then the potential measured as a function of frequency 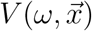 is equal to its variation relative to that of the cell at rest. Thus, for current amplitudes that are not too strong (to remain in the linear regime), 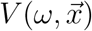 has a very smooth variation in space, despite the fact that the potential at rest may show very abrupt spatial variations near the membrane. In the manuscript, we have designed by equipotential surface any surface for which 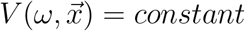 when *ω* = *constant* ≠ 0

## A Equivalent impedance between the extracellular electrode and the ground

In this appendix, we give the explicit expressions to calculate the impedance between the extracellular electrode and the ground in the different experimental conditions considered.

When measuring the equivalent impedance, we have (*Z_a_* || *Z_s_* || *Z_d_*) ⊕ *Z_g_*, where *Z_g_* is the impedance between the ground and the first isopotential surface that surrounds the neuron. *Z_a_* is the impedance of the extracellular medium in contact with the isopotential surface *S*_2_, *Z_s_* is the impedance of the current flowing through the soma membrane in contact with surface *S*_2_, and *Z_d_* is the input impedance of the dendritic tree relative to a reference outside the neuron (Fig. 6).

Thus, we obtain

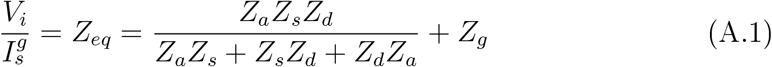

where 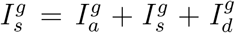. 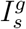 is the generalized current produced by the current source, because in our experiments, the generalized current conservation applies. Note that this does not account for charges created by chemical reactions [36]. We have

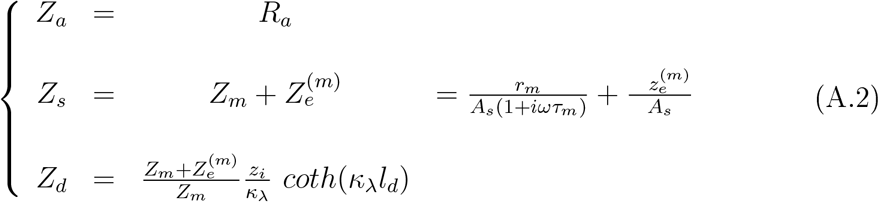

Here, we calculated *Z_d_* as follows. The part of the current source that flows through the dendrite before eventually going to the ground 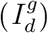 is such that we obtain 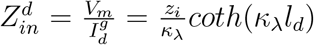 where *V_m_* is the somatic membrane potential at the basis of the dendrite [22]. In addition, applying the generalized current conservation gives the following equality:

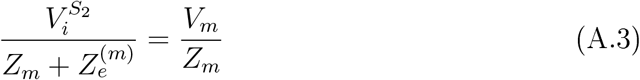

where the potential 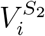 is taken at the isopotential surface *S*_2_. Thus, we have approximately:

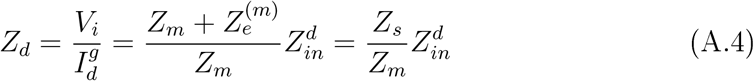

## B RC circuit in series with a resistance

In this appendix, we compare the RC model ((*R* || *C*)) with the RC model in series with a resistance ((*R* || *C*) ⊕ *R*\*). One can see from Fig. B.1 that the impedances of these two models are similar for small frequencies. However, they differ at high frequencies relative to the cut-off frequency of the RC circuit. It is important to also consider that the phase of the RC model tends to −90° when frequency tends to ∞, but it tends to 0° for the RC model in series with a resistance.

**Figure B. 1:**
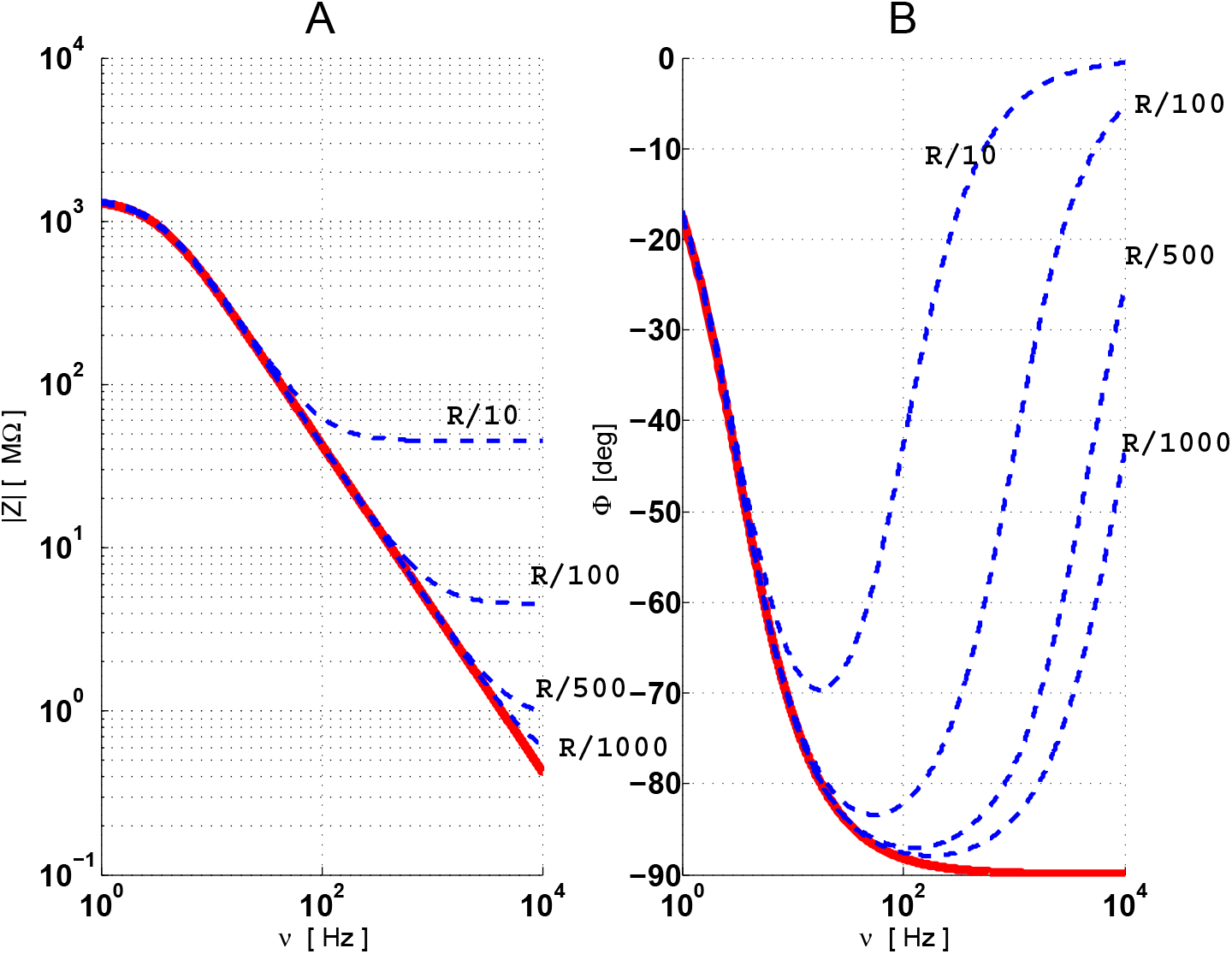
Comparison betwen the impedances of the model RC || (red) and of the model RC || in series with a resistance (blue). The modulus **(A)** and phase **(B)** are shown as a function of frequency.

## C Diffusive impedance in heterogeneous media for a spherical source

In this appendix, we present the theoretical expression of the macroscopic impedance in the case of a diffusive model [20].

In a previous publication [20], we have shown that the macroscopic diffusive impedance (also called Warburg impedance) is derived by a linear approximation of the ratio 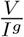, where *V* is the potential difference between the two measurement points. This derivation took into account Boltzmann distribution and Ohm’s law.

The energy given to the charges divides into two parts: one dissipative part (calorific energy) and the part corresponding to the spatial arrangement and distribution of charges as a function of time. The first part is related to Ohm’s law, and the second part to Nernst law.

The presence of a current source in a homogeneous medium breaks its homogeneity. Indeed, the charge distribution around the source cannot be considered constant. The application of Boltzmann’s law in the quasistatic regime (in the sense of classical statistical thermodynamics), in the linear approximation, gives an impedance for a spherical current source of the form:

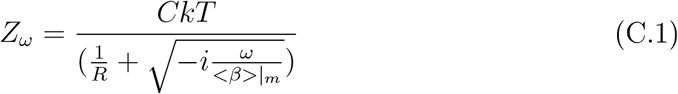

where *C* is a constant which depends on the electric conductivity of the medium in the absence of the source, *R* is the radius of the spherical source (which gives acurvature of 1/*R*^2^), *T* is the absolute temperature in Kelvins, < *β* > |_*m*_ is equivalent to an “effective” diffusion coefficient which is negative, and *k* = 1.38 × 10^-23^ *J*/°*K* isthe Boltzmann constant (for more details, see [20]). This model is called “diffusive model”, and is used here for the particular case of a spherical source.

By setting

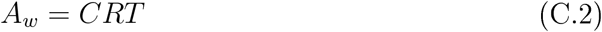

and

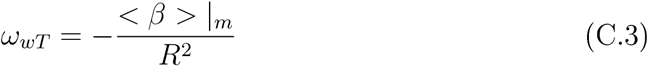

we can write expression C. 1 as above:

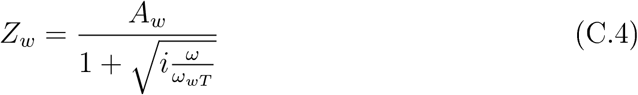

At constant temperature, the measurement of the impedance allows one to determine the values of *A_w_* and *ω_wT_* = 2*πν_wT_*. The parameters *A_w_* and *ω_wT_* are real.

However, we assumed

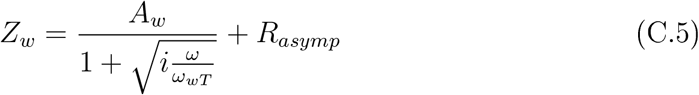

because the original derivation of the expression of the Warburg impedance in mean-field [20] considered the particular solution of the differential equations in mean-field, also called the “forced solution”. The general solution is the sum of this particular solution and the solution of the homogeneous equation (▿^2^*V_ω_* = 0). To take this into account, one needs to add a resistance in series with the forced solution (*R_asymp_*.). This asymptotic resistance appears at very large frequencies in the experimental measurements (see Results).

**Figure D. 1:**
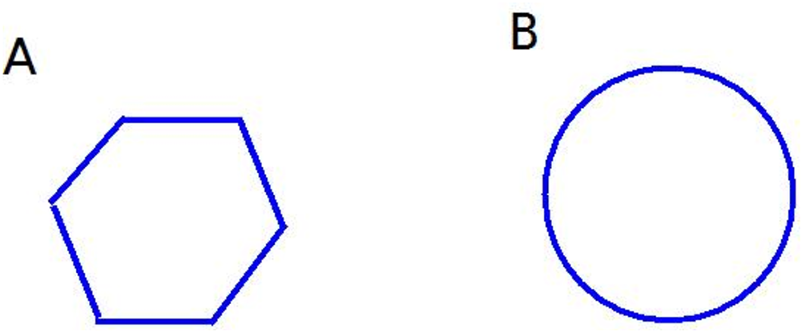
Examples of how the curvature of isopotential surfaces determines the cutoff frequency of the impedance. A. Volume delimited by a plane (infinite curvature radius), resulting in a cutoff frequency near zero. B. Similar volume delimited by a border of constant curvature. In this case, the cutoff frequency is larger because it is inversely proportional to the curvature radius (Appendix D).

## D Threshold frequency and surface curvature of the diffusive model in the general case

In this appendix, we give some details about the relation between the threshold frequency in the diffusive impedance (expression C.5) and the curvature at a given point of a surface *S*. In other words, we show how to apply the diffusive model to surfaces that are non-spherical.

The diffusive model of Appendix C can be applied to an arbitrary surface because we can build an approximately continuous surface 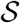 by the sum of portions of spherical surfaces centered on different points of 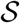, where the curvature corresponds to that of 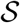. Note that the smaller the intrinsic curvature of a surface, the smaller is the threshold frequency of that surface^10^.

For example, if we have a surface 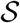 composed of two spherical portions (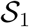 and 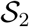) of very different radius, the diffusive impedance diffusive as sensed by the surface is equal to the two impedances of each portion in parallel, because the current divides between both of them. It follows that *Z_S_* = *Z*_*S*_1__ || *Z*_*S*_2__, with:

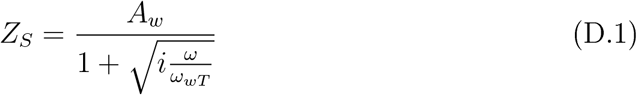

where

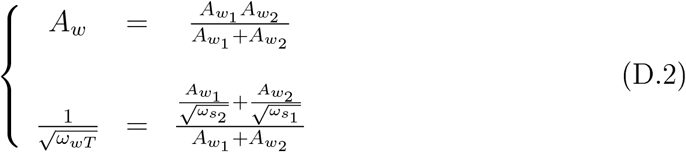

If the surfaces 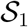 and 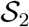 have the same impedance, then the impedance of 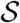 is twice smaller, but the threshold frequency remains the same. If each surface displays a similar Warburg amplitude but with different threshold frequencies, then we obtain

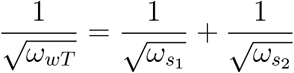

It follows that if we approximate a given surface with a set of *N* spherical portions of same Warburg amplitude, the threshold frequency is given by:

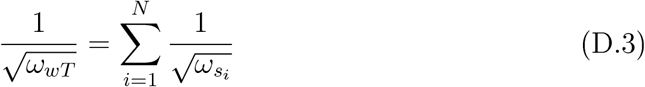

Thus, the portions of surface with the smallest curvature will determine the threshold frequency of the ensemble, Consequently, it is possible to obtain a very small threshold frequency, even in a domain of a very small volume (Fig. D.1).

## E Macroscopic impedance relative to ground

In this appendix, we consider the macroscopic impedance as sensed by the electrode injecting the current in the soma, via the dendrite, before reaching the ground, 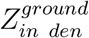, and the impedance as seen by the current going to the ground independently of the dendrite, 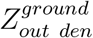, for a ball-and-stick model in a resistive extracellular medium. Note that the impedance between the soma and the ground is given by 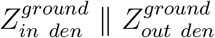.

We also consider that the cytoplasm is resistive, as well as the Debye layers surrounding the membrane. We will consider the experimental measurements of Section 3.2. Importantly, in the present experiments, the impedance between the cell and the ground should be calculated in an “open” configuration, because the current injected in the neuron flows to the ground without looping back to the neuron.

We numerically compared the impact of the two configurations, open and closed, at the basis of the dendrite (stick). In particular, the parameter *k*_λ_ = *k*/λ is a good indicator to evaluate the differences between the two configurations.

For this purpose, we first assumed that the extracellular impedance 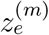 has the same value at every point in the membrane in soma and dendrites. This hypothesis is reasonable if the neuron is physically smaller than the geometrical dimensions of the experimental preparation. Indeed, this parameter measures the impedance of the extracellular medium as sensed by the membrane, as defined by 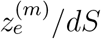 which is the impedance between *dS*, a differential element of membrane, and the ground.

**Figure E.1:**
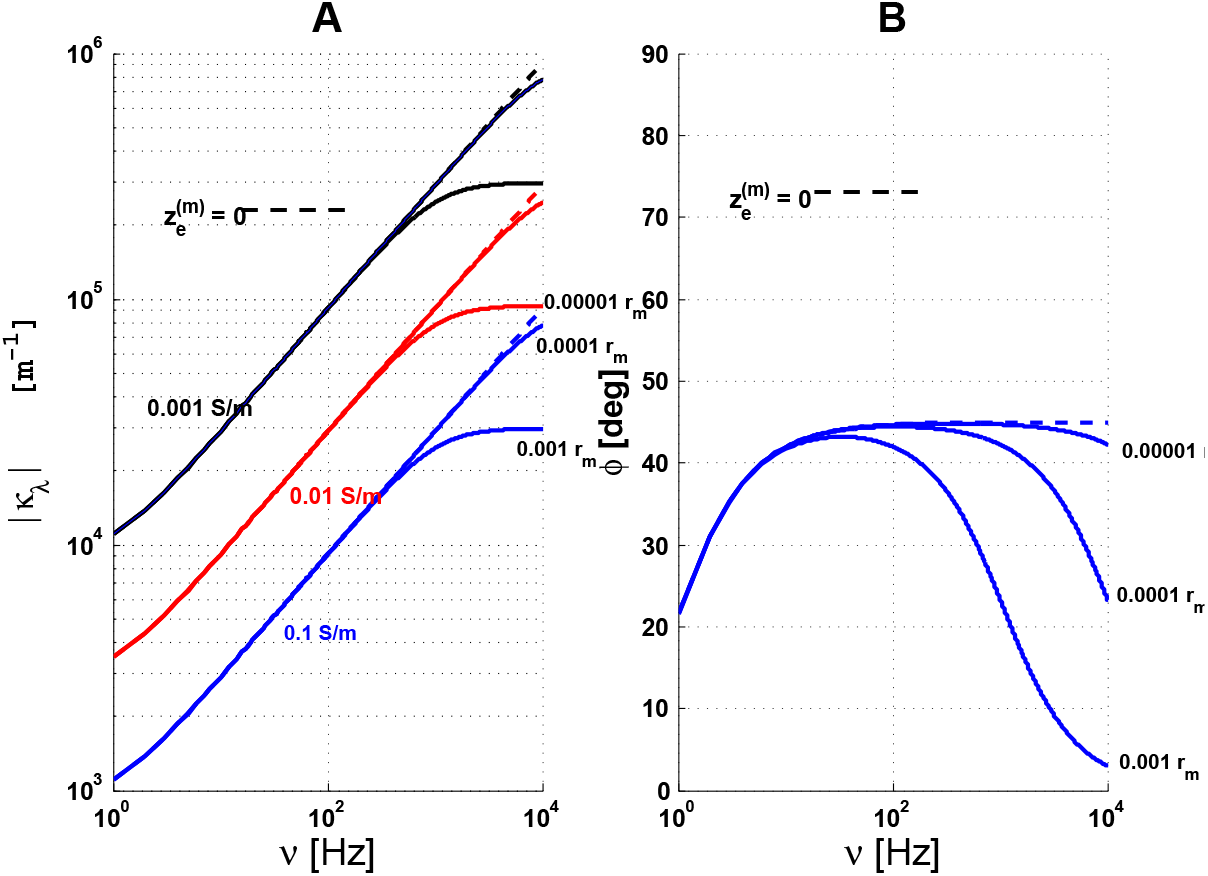
Graph of 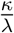 as a function of frequency for the ball-and-stick model. Here, 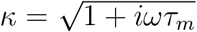, the soma has a radius of 10 *μm*, while the length and diameter of the stick are respectively of 600 *μm* and 3 *μm*. In this example, the membrane time constant *τ_m_* is of 30 *ms*, *c_m_* = 0.01 *F*/*m*^2^ is the specific membrane capacitance, and 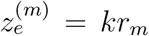 with *k* = 0.001, 0.0001, 0.00001. The dashed lines correspond to the open configuration, and continuous lines to the closed configuration. The electric conductivity of the cytoplasm 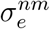 corresponds to the different colors, Blue: 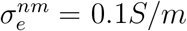, Red: 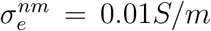, Black: 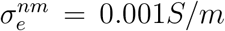. Note that in the case 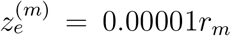 is approximately equivalent to a supraconductive medium, for frequencies smaller than 10 *kHz*.

According to the generalized cable theory [22], for a resistive medium, we have:

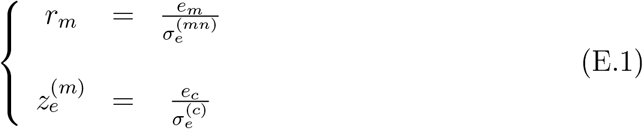

where *e_m_* and *e_c_* are the thickness of the membrane and of Debye layers, respectively. 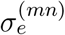 and 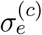 are the mean electric conductivity of the membrane and of the extracellular medium (comprising Debye layers), respectively. Debye layers have a high density of ions, and thus have a different conductivity than the “bulk” of the medium. The ions around the membrane are distributed according to Boltzmann distribution, forming Debye layers, and the diffusive model must be taken into account in this case (Appendices C and D). The electric conductivity is much lower in Debye layers compared to the other parts of the extracellular medium, which is considered homogeneous away of Debye layers. We assumed a thickness equivalent to that of Debye layers in the expression of 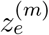 (Eq. E.1).

However, in this appendix, we neglect the possible frequency dependence and model the impedance of Debye layers with a resistance, as if the threshold frequency was very large. The goal here is to determine, as simple as possible, the physical consequences of the magnitude of 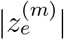 relative to *r_m_* on the current division between the soma and the dendritic stick.

According to expressions E.1, we obtain:

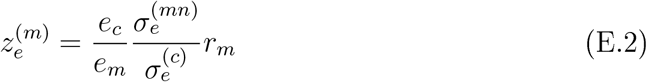

where *r_m_* = *τ_m_*/*c_m_* = 100*τ_m_* (with *c_m_* = 0.01 *F*/*m*^2^).

For a value of *τ_m_* = 30 *ms*, *e_c_* = 0.1*e_m_* and 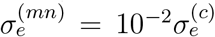, we obtain 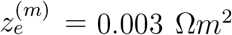. This value gives the order of magnitude of the physical effects on the impedances 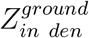 and 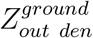. The value of the membrane time constant is that of the experiments presented here.

Next, from the evolution of the electric conductivity of the cytoplasm, we consider three different values: 0.1, 0.01 and 0.001 S/m (Fig. E.2) The first value approximately corresponds to that of ACSF for a temperature of 37°*C*. The two other values are smaller, to simulate the fact that the cytoplasm is a heterogeneous medium (presence of organites), which creates a tortuosity, as well as electric polarization. These effects have been reported to diminish electric conductivity [27, 28].

**Figure E.2:**
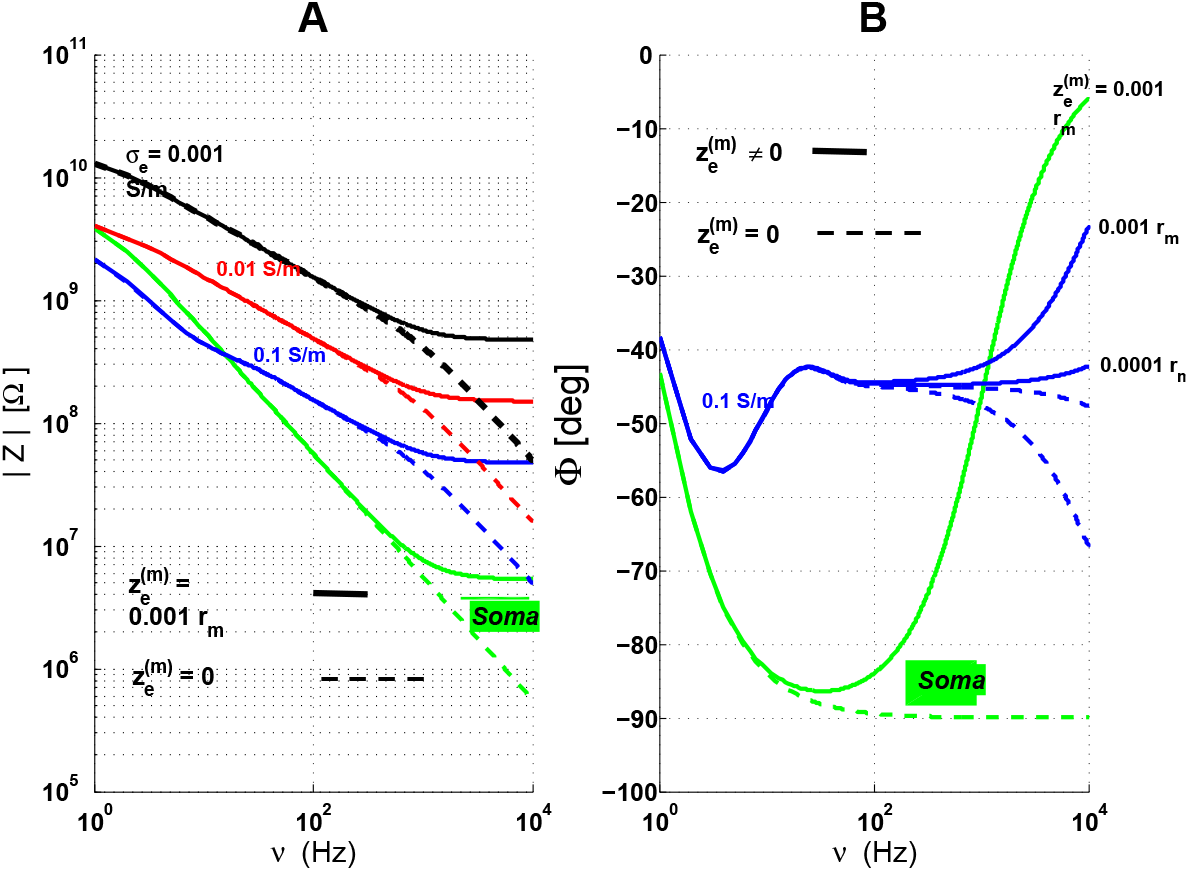
Example of input impedance 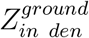 and of soma impedance 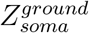. Here, 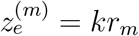 and *k* = 0.001, 0.0001. The electric conductivity of the cytoplasm is 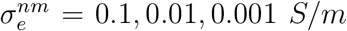. The other parameters are the same as in Fig. D.1. The dotted lines correspond to the case 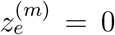, which is equivalent to neglect Debye layers; solid lines correspond to a resistive model, with Debye layers taken into account.

Figure E.2 shows examples of the input impedance in the following conditions. **1)** The open and closed configurations give very different results when 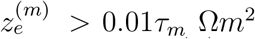, otherwise the differences are small for parameters *κ*_λ_. Note that the case 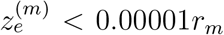 is as if Debye layers were inexistent for frequencies smaller than 10 *kHz*. **2).** For 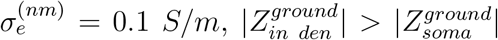 if *ν* > 100 *Hz*, for 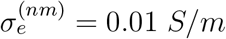. This inequality holds up to about 1 Hz. For 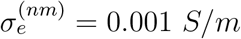, the modulus of the dendrite (stick) impedance is much larger than that of the soma.

## F Apparent electric conductivity and permittivity

In this appendix, we define the apparent electric conductivity and permittivity. In general, we have the following linking relations between the diffusion 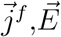 and the 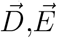:

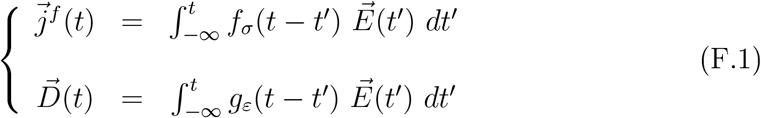

This is obtained in the framework of a mean-held theory of Maxwell equations when the extracellular and intracellular media are linear and homogeneous [20, 35]. Note that the functions *f_σ_* and *g_ε_* are real functions which can model different physical phenomena, such as ionic diffusion, electric polarization, calorific (resistive) dissipation, etc. The integral expresses the fact that the free-charge current density and displacement current density at a given time *t* are not only determined by the electric field at time *t* but also by the whole history of its time variations. These functions are the inverse Fourier transform of electric conductivity and permittivity expressed in Fourier frequency space^11^. For example, for an ideal electric resistance, we have *f_σ_*(*t*) = *σ_e_δ*(*t*), for an ideal capacitance we have *g_ε_*(*t*) = *ε_s_δ*(*t*) (where *σ_e_* are *ε_s_* are constant in time). These two parameters are respectively the electric conductivity and permittivity. In these two ideal cases, the relations F. 1 give the following equalities: 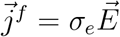 and 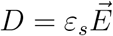, where *σ_e_* and *ε_s_* are time independent. Note that these two ideal elements have no memory of the past (which is expressed by the Dirac deltas), and this is not generally the case of frequency-dependent electric conductivity and permittivity.

Consequently, we have in general, in Fourier frequency space:

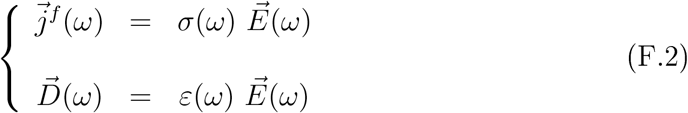

where *σ*(*ω*) and *ε*(*ω*) are respectively the Fourier transforms of *f_σ_*(*t*) and *g_ε_*(*t*) Because *f_σ_*(*t*) and *g_ε_*(*t*) are real functions, this implies in general *σ*(–*ω*) = *σ*\*(*ω*) and *ε*(–*ω*) = *ε*\*(*ω*). Here, the real parts are necessarily even functions and the imaginary parts are odd functions. The relations F.2 imply that the generalized current density is related to the electric field by:

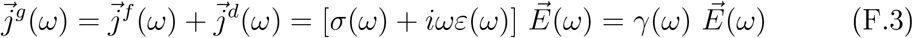

where *σ* = *σ*′ + *iσ*″ and *ε* = *ε*′ + *iε*″ are complex functions in general, while *σ*′, *σ*″, *ε*′, *ε*″ are real functions. We have the following particular cases: an ideal resistance is such that we have *σ*(*ω*) = *σ_e_* (Ohm’s law), an ideal capacitance corresponds to *ε*(*ω*) = *ε*, so that *σ* and *ε* are real numbers and do not depend on frequency.

From these relations, we obtain:

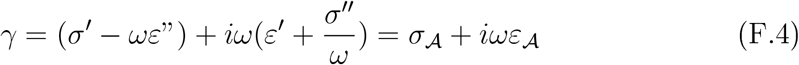

which defines the apparent electric conductivity 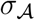 and the apparent electric permittivity 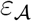. In general, the apparent electric permittivity can be viewed as a type of resistance which depends on frequency and allows to calculate the dissipated power at a given frequency. The ratio 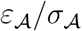 can be used to evaluate the relaxation time of the medium. Note that this definition corresponds to the electric parameters measured in previous studies [7, 34]. In these experimental studies, the measurements are characterized by parameters 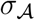 and 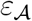 because we interpret the experimental measurements in a very heterogeneous medium as if it was a non-ideal resistance (which depends on frequency) in parallel with a non-ideal capacitance (which also depends on frequency). A heterogeneous medium can be modeled as a homogeneous medium where the parameters depend on frequency with respect to macroscopic measurements. This is analogous to classical thermodynamics where pressure and temperature can be used to characterize a physical system.

Note that the apparent electric permittivity tends to infinity if the imaginary part of the electric conductivity does not tend to zero at null frequency. This is not the case for an ideal resistance because the imaginary part of its electric conductivity is zero. However, for a diffusive (planar) impedance (with zero curvature, see Appendix D) the imaginary part of electric conductivity is non-zero, since in this case 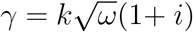 where *k* is a constant.

By definition, the complex admittance *Y* between the two arms of a plane capacitor with a given medium in between, is given by 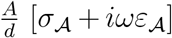. *A* is the arms area and *d* is the distance separating them. If we assume that these geometrical dimensions do not generate boundary effects, the electric field between the arms is of *V*/*d* where *V* is the voltage difference between the arms of the capacitor.

**Figure F.1:**
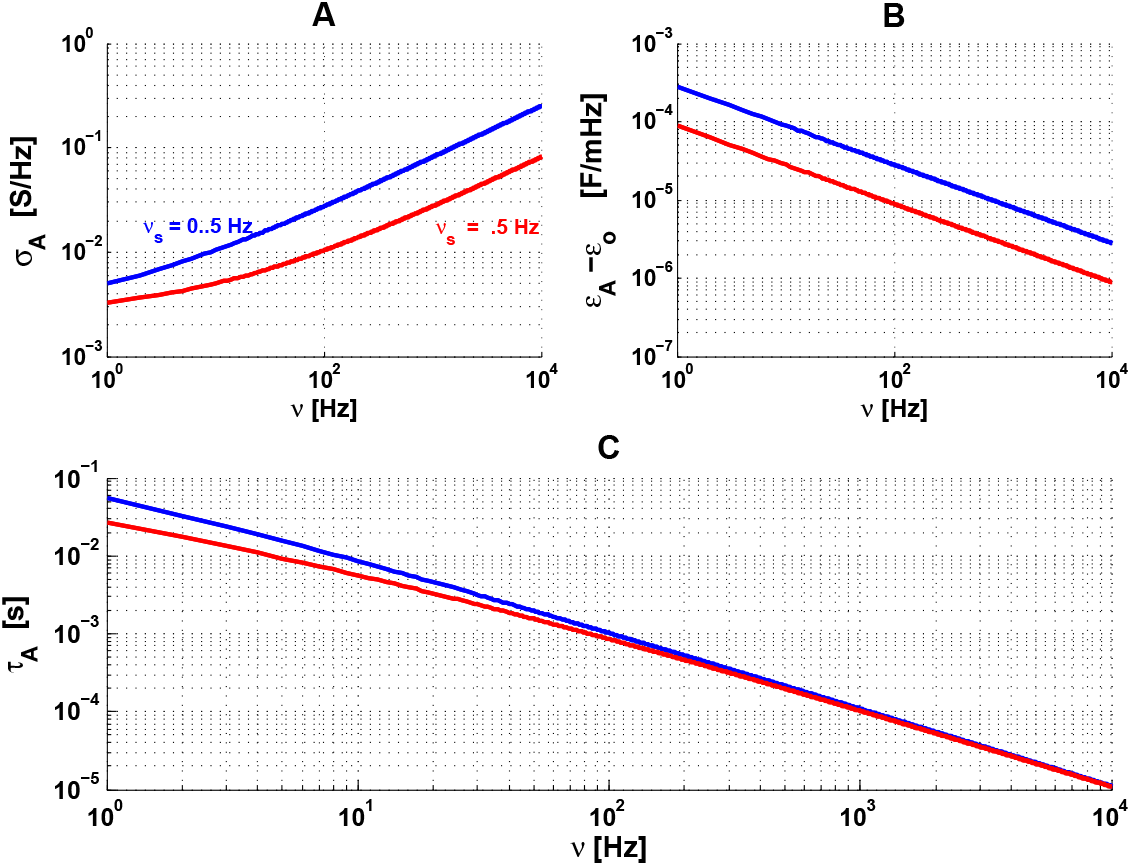
Apparent conductivity and permittivity of a diffusive impedance. The red and blue curves correspond to *ν_wT_* = .5 *Hz* and black curves correspond to *ν_wT_* = 40 *Hz*. We have *A_w_* = 16 *M*Ω and *A*/*d* = 10 *μm* for all curves,

For example, the measurement of the apparent parameters of a medium with a diffusive impedance gives the following equality:

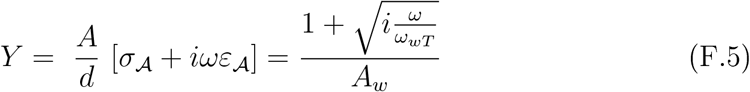

It follows that the frequency dependence of the parameters is given by the following expressions:

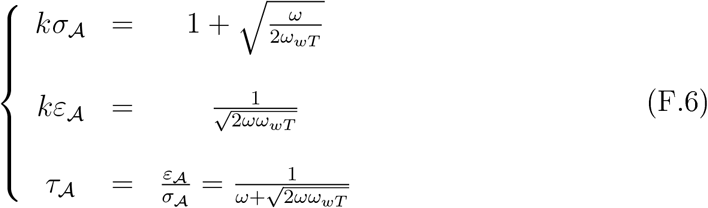

where the constant *k* is equal to 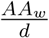.

Thus, the apparent electric conductivity tends to *k*, the electric permittivity tends to infinite, and the dielectric relaxation time tends to infinity for *ω* → 0. We conclude that if ionic diffusion is not negligible, then the linear approximation of its effect on the measured impedance is as if the dielectric relaxation time tends to infinity at null frequency (Fig. F.1).

## G Ensemble of the measurements

In this appendix, we show the ensemble of experimental results obtained in the different preparations. Figures G.1 (non-arborized neurons in culture), G.3 (arborized neurons in culture) and G.5 (arborized neurons in brain slices) respectively show the impedance of Region 1-2 (between the intracellular and extracellular electrodes). The same preparations are respectively shown in Figs. G.2, G.4 and G.6 for the modulus of the impedance of Region 2-3 (between the extracellular electrode and the ground). The values of the experimental parameters for the different experimental preparations are shown respectively in Table G.1, G.3 and G.5, while the values of the corresponding models are shown in Table G.2, G.4 and G.6.

**Figure G.1:**
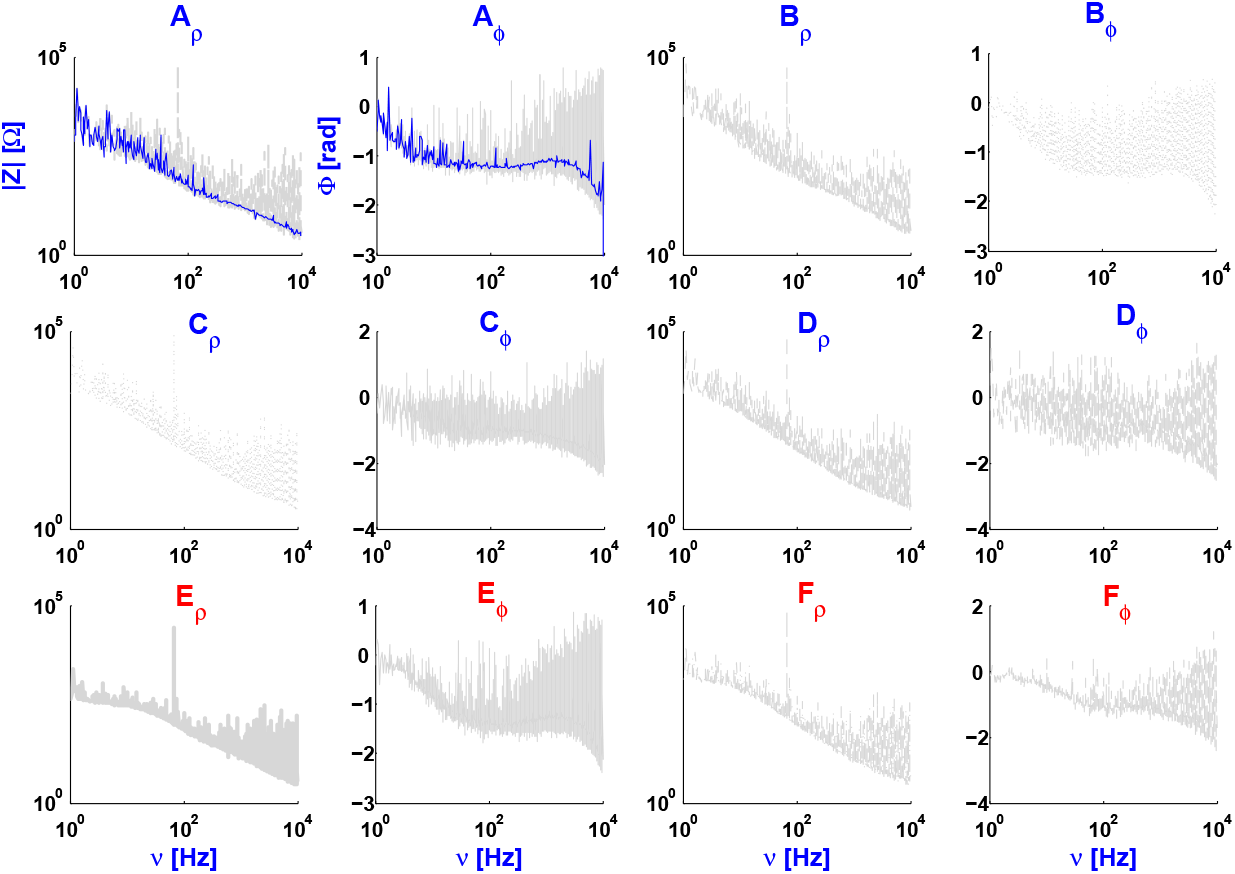
Experimental measurement between intracellular and extracellular electrodes for 6 non-arborized neurons in culture. On the basis of the fits, two groups can be distinguished, one with *τ_m_* around 30 ms and another group with 5-15 ms (Tables G.1 and G.2). The blue curves are cubic spline fits of the experimental data.

**Figure G.2:**
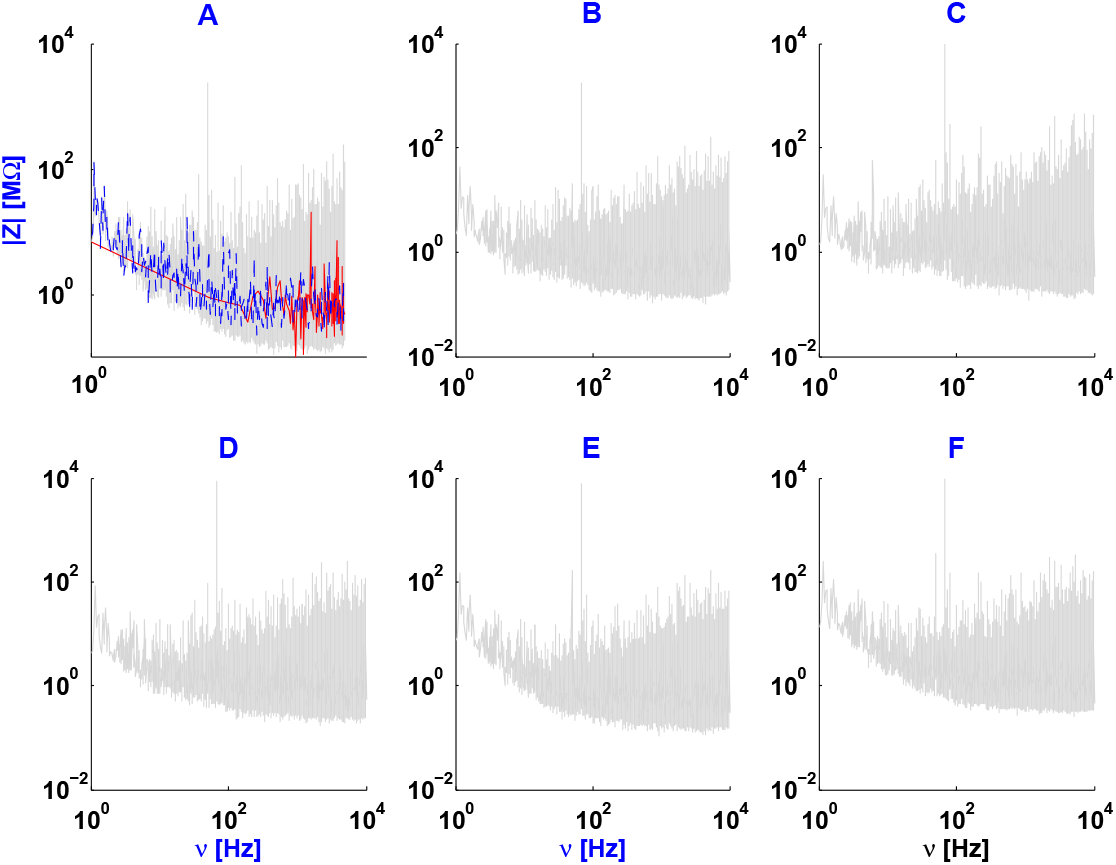
Experimental measurements between extracellular and ground for 6 non-arborized neurons in primary cell culture, as shown in Tables G.1 and G.2. The blue curves are cubic spline fits to the logarithm of the experimental data, while the red curves are the direct cubic spline fits of the data.

**Table G.1:** Individual experimental parameters for 6 non-arborized neurons in culture, shown in Fig. G.1. In the absence of measurements, a NaN is indicated.

| | $V_m$ (mean ± standard deviation) during white noise injection (mV) | $\sigma$ of white noise current (pA) | Number of sweeps | Membrane resistance ($M\Omega$)<br>Membrane time constant (ms) |
| --- | --- | --- | --- | --- |
| **A** : 2020021702 | $-68.9 \pm 2.5$ | 10 | 18 | NaN<br>NaN |
| **B** : 2020021801 | $-64.6 \pm 6.4$ | 5 | 13 | 2995<br>117 |
| **C** : 2020021901 | $-62.3 \pm 2.5$ | 10 | 20 | 900<br>62 |
| **D** : 2020021803 | $-57.7 \pm 4.0$ | 5 | 71 | 2090<br>105 |
| **E** : 2020021802 | $-64.3 \pm 1.6$ | 10 | 14 | 558<br>16 |
| **F** : 2020031503 | $-57.1 \pm .9$ | 5 | 25 | 1339<br>32 |

**Table G.2:** Parameters for the diffusive model for each non-arborized neuron in Table G.1. 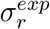 and 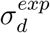 are respectively the mean square error of resistive and diffusive models relative to the experimental measurements for each neuron.

| | $R_m$<br>[$M\Omega$] | $\tau_m$<br>[ms] | $A_w$<br>[$M\Omega$] | $\nu_{wT}$<br>[ms] | $R_{asympt}$<br>[$M\Omega$] | $\sigma_d^{exp}$<br>[$M\Omega$] | $\sigma_r^{exp}$<br>[$M\Omega$] |
| --- | --- | --- | --- | --- | --- | --- | --- |
| **A** | 1000 | 30 | 625 | 0.1 | 4 | 3.4 | 142.9 |
| **B** | 3000 | 30 | 2000 | 0.2 | 4 | 16 | 225.0 |
| **C** | 3000 | 35 | 2000 | 0.05 | 6 | 3.5 | 210.0 |
| **D** | 3000 | 25 | 2000 | 0.05 | 3 | 3.8 | 215.0 |
| **E** | 3000 | 5 | 2000 | 0.7 | 2 | 2.9 | 58.3 |
| **F** | 1000 | 15 | 670 | 0,1 | 2 | 2.2 | 179.0 |

**Figure G.3:**
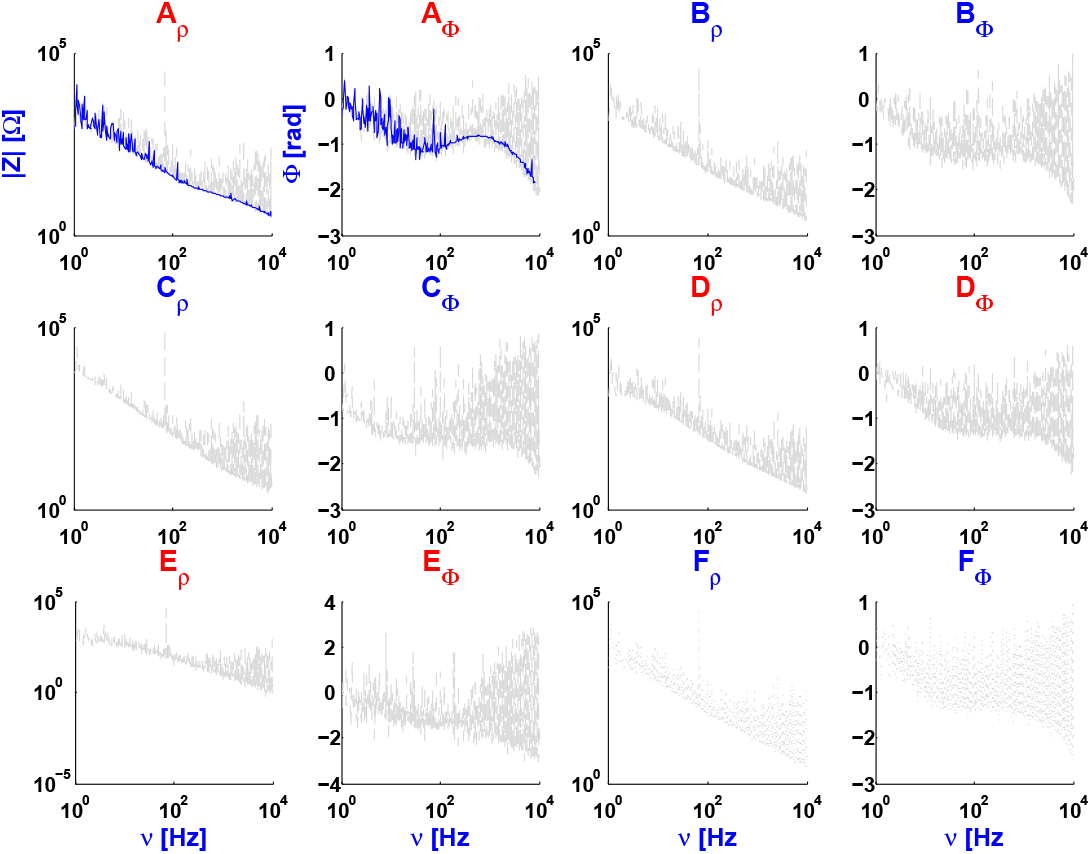
Experimental measurements between intracellular and extracellular electrodes for 6 arborized neurons in primary cell culture (quasi-homogeneous medium). The blue curves are cubic spline fits of the experimental data.

**Figure G.4:**
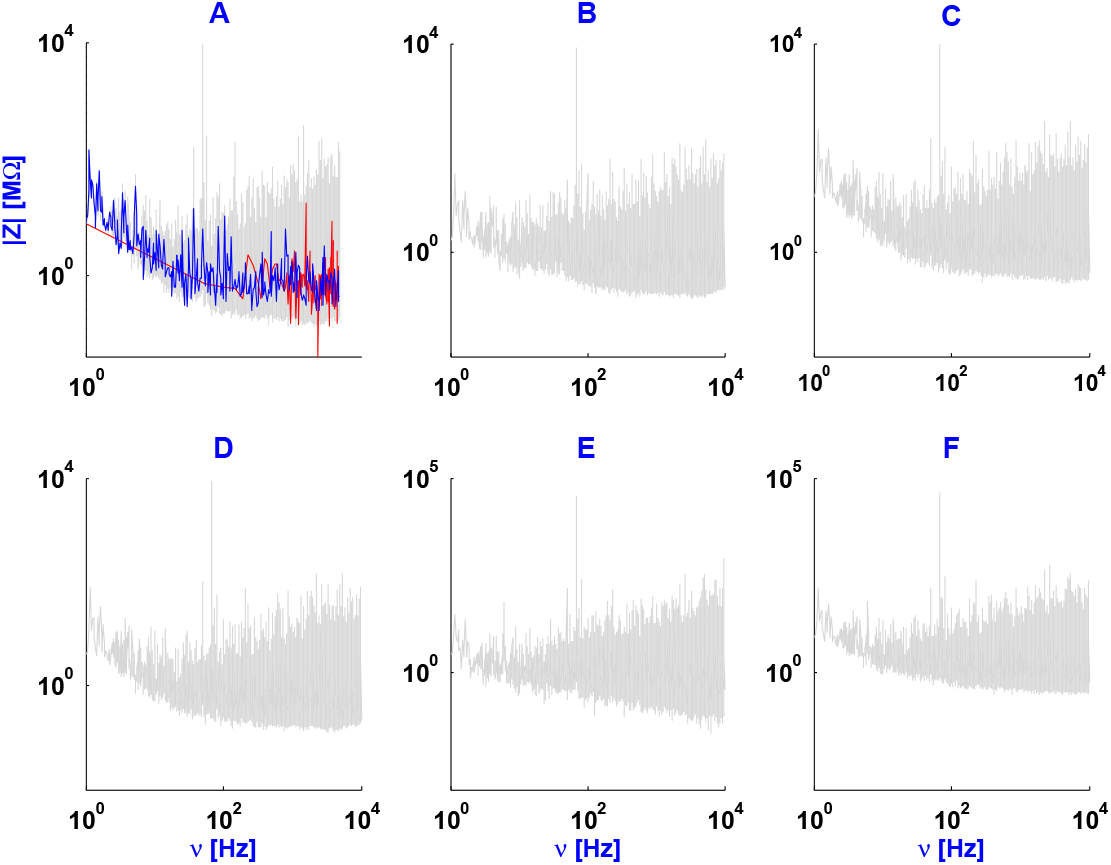
Experimental measurements between extra and ground for 6 arborized neurons in culture, as shown in Tables G.3 and G.4. The blue curves are cubic spline fits to the logarithm of the experimental data, while the red curves are the direct cubic spline fits of the data.

**Table G.3:** Individual experimental parameters for 6 arborized neurons in culture.

| | $V_m$ (mean ±<br>standard deviation)<br>during white<br>noise injection (mV) | $\sigma$ of white<br>noise<br>current<br>(pA) | Number<br>of sweeps | Membrane<br>resistance ($M\Omega$)<br>Membrane time<br>constant (ms) |
| --- | --- | --- | --- | --- |
| **A** : 2020021701 | $-66.0 \pm 2.6$ | 10 | 46 | 549<br>25 |
| **B** : 2020021902 | $-68.1 \pm 4.0$ | 10 | 31 | 1820<br>67 |
| **C** : 2020031501 | $-76.1 \pm 4.4$ | 5 | 28 | 4737<br>109 |
| **D** : 2020031502 | $-67.5 \pm 4.4$ | 10 | 60 | 1347<br>31 |
| **E** : 2020031602 | $-66.7 \pm 2.1$ | 10 | 3 | 793<br>22 |
| **F** : 2020031603 | $-69.4 \pm 2.7$ | 5 | 26 | 2220<br>89 |

**Table G.4:**
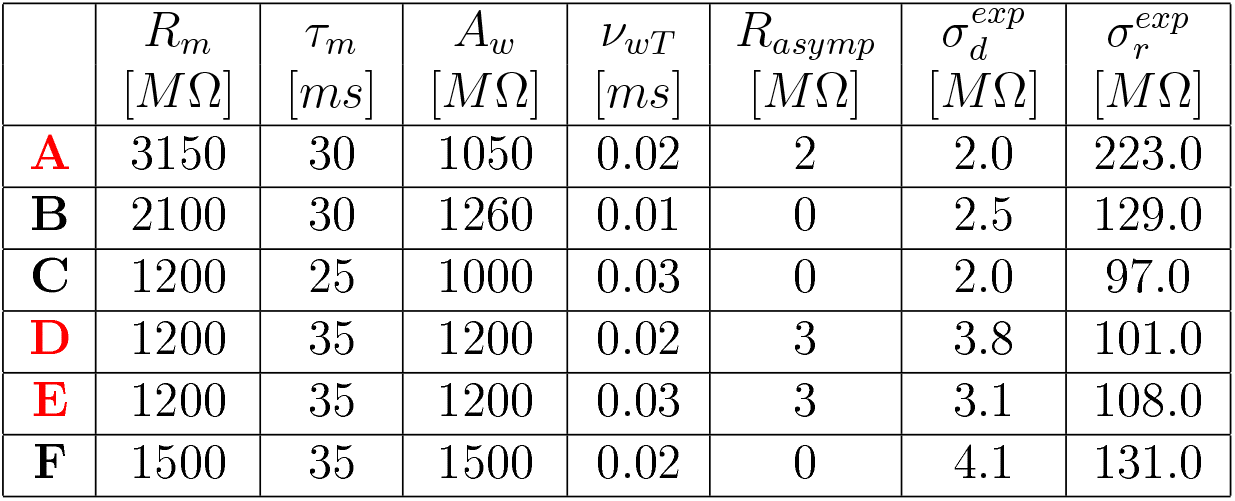
Individual parameters obtained from fits for 6 arborized neurons in culture from Table G.3. 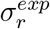 and 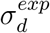 are respectively the mean square error of resistive and diffusive models with respect to the experimental data.

**Figure G5:**
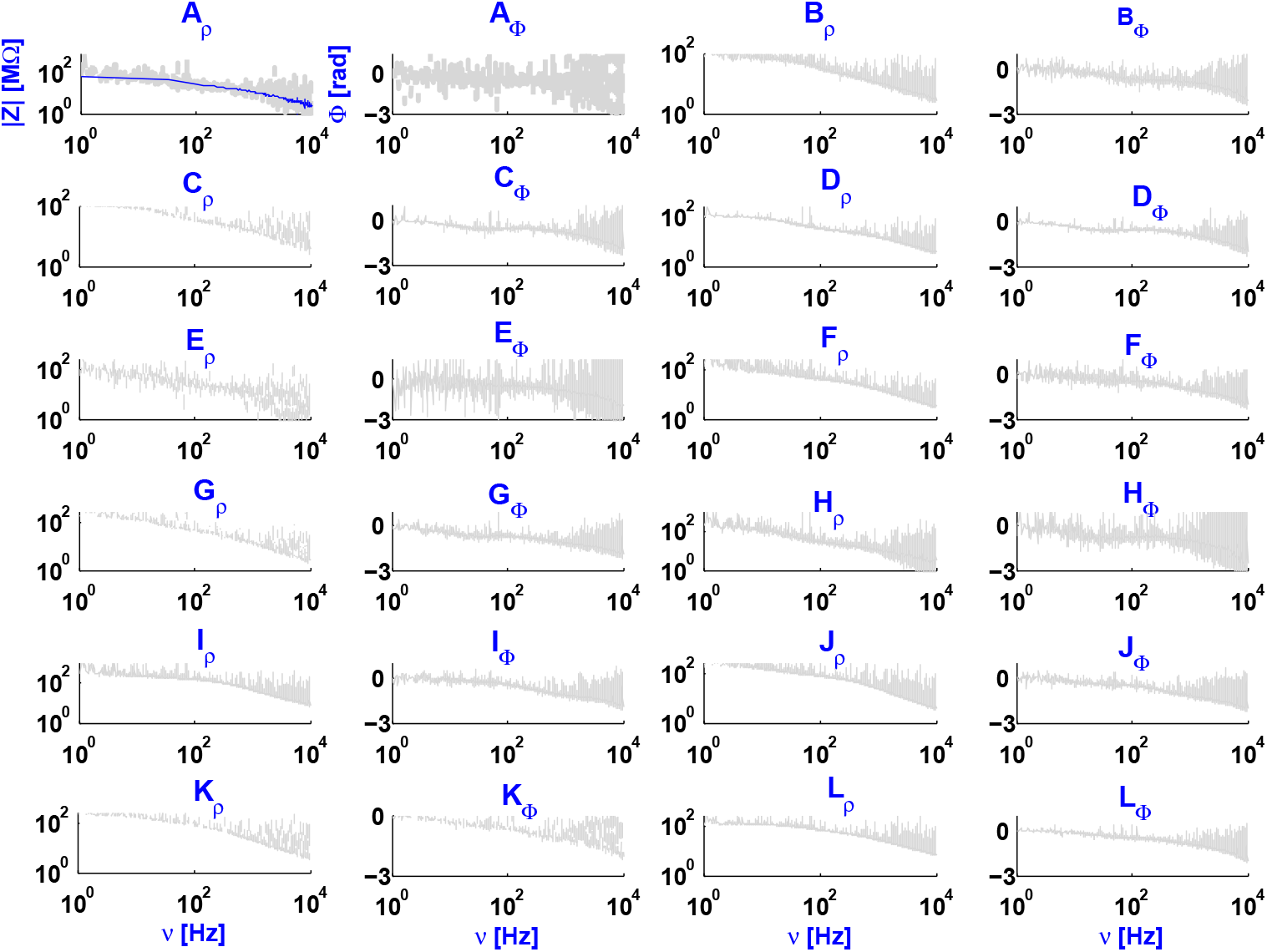
Experimental measurements between intracellular and extracellular electrodes for 9 arborized neurons in brain slices, as shown in Tables G.5 and G.6. The blue curves are the cubic spline fits of the experimental data.

**Figure G.6:**
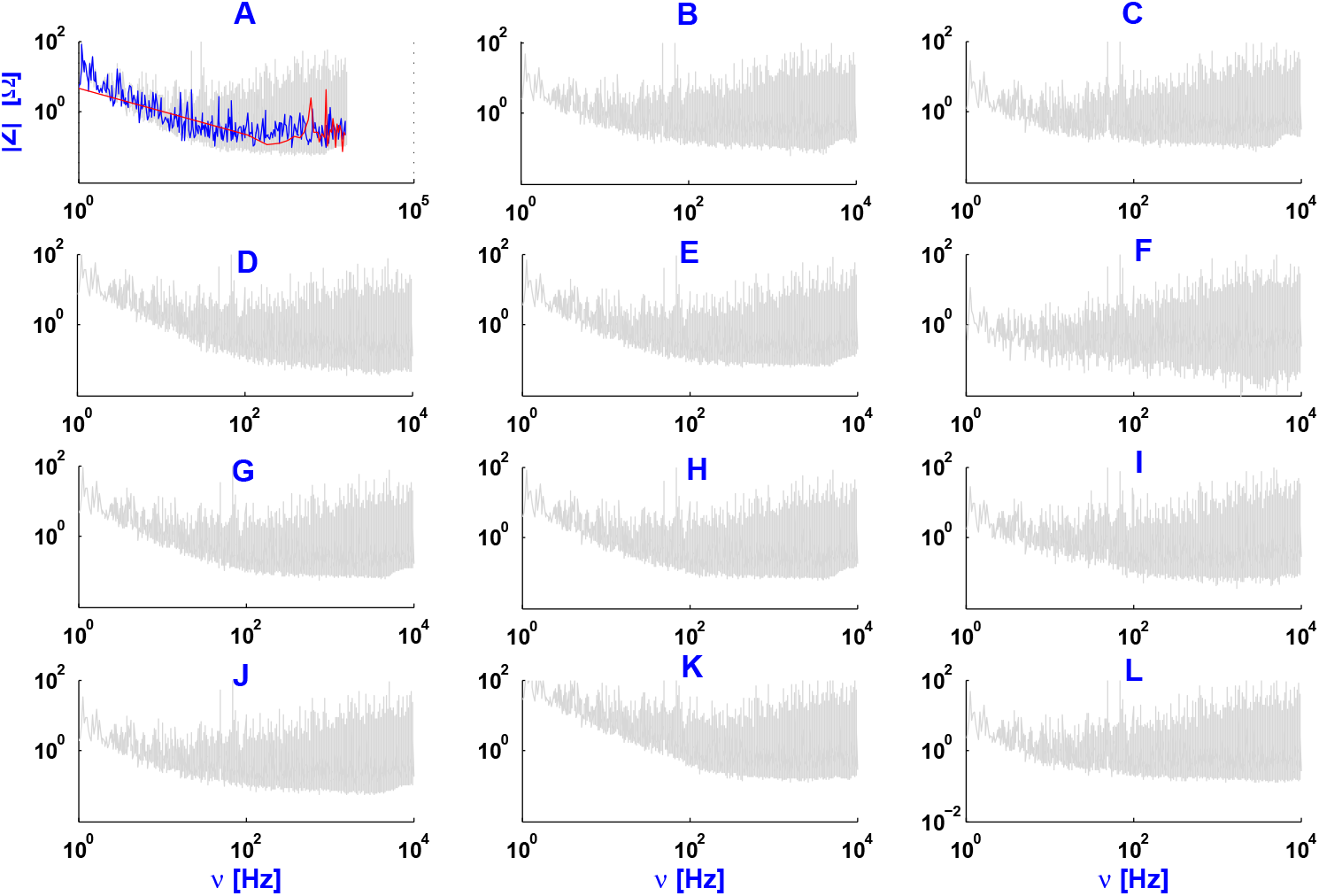
Experimental measurements between extracellular and ground for arborized neurons in slices, as shown in Tables G.5 and G.6. The blue curves are cubic spline fits to the logarithm of the experimental data, and the red curves are the direct cubic spline fits of the data.

**Table G.5:** Individual experimental parameters for 9 arborized neurons in brain slices (MSN: medium-sized spiny neuron; FS: fast-spiking interneuron).

| | $V_m$ (mean ±<br>standard deviation)<br>during white<br>noise injection (mV) | $\sigma$ of white<br>noise<br>current<br>(pA) | Number<br>of sweeps | Membrane<br>resistance (MΩ)<br>Membrane time<br>constant (ms) |
| --- | --- | --- | --- | --- |
| **A** : 2018121204 (MSN) | $-75.2 \pm 1.1$ | 20 | 1 | 67<br>6.0 |
| **B** : 2018121205 (FS) | $-74.8 \pm 1.2$ | 20 | 10 | 53<br>5.5 |
| **C** : 2018122701 (FS) | $-78.2 \pm 1.3$ | 20 | 15 | 124<br>5.6 |
| **D** : 2019021202 (MSN) | $-75.8 \pm 1.3$ | 20 | 6 | 83<br>4.2 |
| **E** : 2019021203 (MSN) | $-73.9 \pm 1.7$ | 20 | 59 | 85<br>4.0 |
| **F** : 2019021302* (MSN) | $-69.1 \pm 1.8$ | 20 | 13 | 64<br>7.7 |
| **G** : 2019021304* (MSN) | $-69.3 \pm 1.7$ | 20 | 10 | 128<br>9.5 |
| **H** : 2019021401 (MSN) | $-64.6 \pm 1.4$ | 10 | 5 | 124<br>8.4 |
| **I** : 2019021801 (MSN) | $-80.1 \pm 2.7$ | 10 | 31 | 92<br>2.8 |
| **J** : 2019021802<br>(Cholinergic interneuron) | $-73.3 \pm 1.6$ | 10 | 35 | 158<br>6.3 |
| **K** : 2019021803* (MSN) | $-67.1 \pm 1.6$ | 10 | 8 | 206<br>7.9 |
| **L** : 2019021901 (MSN) | $-76.8 \pm 1.4$ | 10 | 49 | 44<br>3.4 |

**Table G.6:** Individual parameters obtained from the fits for arborized neurons in brain slices from Table G.5. The total resistance for *ν* = 0 is equal to 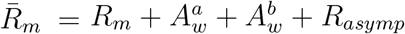. 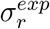 and 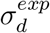 are the mean square errors of the resistive and diffusive models relative to experimental data, respectively.

| | $\bar{R}_m$ | $R_m$ | $\tau_m$ | $A_w^a$ | $\nu_{wT}^a$ | $A_w^b$ | $\nu_{wT}^b$ | $R_{asympt}$ | $\sigma_d^{exp}$ | $\sigma_r^{exp}$ |
| --- | --- | --- | --- | --- | --- | --- | --- | --- | --- | --- |
| | [$M\Omega$] | [$M\Omega$] | [$ms$] | [$M\Omega$] | [$Hz$] | [$M\Omega$] | [$Hz$] | [$M\Omega$] | [$M\Omega$] | [$M\Omega$] |
| **A** | 120 | 48 | 5 | 48 | 5 | 24 | 40 | 1 | 2.1 | 6.0 |
| **B** | 120 | 50 | 5 | 50 | 5 | 24 | 40 | 4 | 2.3 | 6.6 |
| **C** | 140 | 47 | 5 | 47 | 5 | 40 | 60 | 4 | 2.3 | 5.2 |
| **D** | 140 | 48 | 5 | 48 | 1 | 40 | 60 | 2 | 2.4 | 5.1 |
| **E** | 140 | 48 | 2.5 | 48 | 1 | 40 | 60 | 1 | 2.0 | 6.0 |
| **F** | 140 | 48 | 5 | 48 | 1 | 40 | 60 | 4 | 5.5 | 11.0 |
| **G** | 256 | 90 | 20 | 90 | 1 | 75 | 40 | 1 | 6.3 | 9.5 |
| **H** | 256 | 90 | 20 | 90 | 0.1 | 75 | 10 | 1 | 3.0 | 10.1 |
| **I** | 426 | 150 | 1 | 150 | 1 | 125 | 20 | 1 | 28.1 | 72.0 |
| **J** | 320.5 | 90 | 10 | 90 | 20 | 125 | 60 | .5 | 6.1 | 26.3 |
| **K** | 320.5 | 90 | 10 | 90 | 20 | 75 | 60 | .5 | 6.1 | 26.3 |
| **L** | 320.5 | 48 | 1 | 48 | 45 | 40 | 80 | 3 | 6.3 | 15.3 |

The extracellular-to-ground measurements are similar in all preparations and exhibit a similar frequency dependence. All fits show that the impedance modulus of the extracellular medium is of the order of that of ACSF.

However, this is not the case for the intracellular-to-extracellular measurements in the different preparations. For all cells and for all preparations, the diffusive model fits better the experimental data.

Finally, for the different preparations, the experimental measurements (membrane resistance and membrane time constant) shown in Table G.1 are different from that displayed in Table G.2. This difference shows that the evaluation of the membrane time constant using a current pulse leads to different membrane time constants as those evaluated from direct fitting of the impedance. This aspect is examined in more detail in the following (Appendix H).

## H Voltage between two points for a square current pulse

In this appendix, we calculate the voltage between two points when the injected current is a square pulse.

We model a pulse of current as follows:

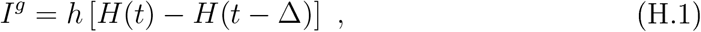

where Δ > 0 is duration of the current pulse, and *H* is the Heaviside function. The most general linear relation between current and voltage is given by the following expression:

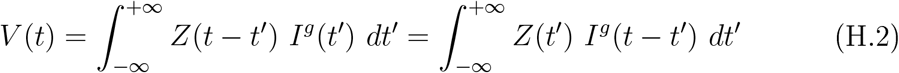

It follows that the derivative of the voltage is given by:

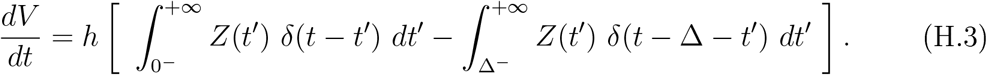

with 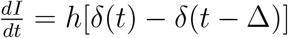.

Thus, we obtain the following equality:

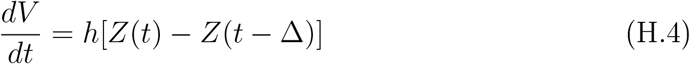

According to the complex Fourier transform,

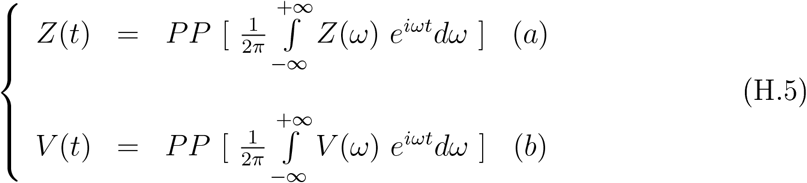

where PP means the principal part of the integral.

In a diffusive medium, we have:

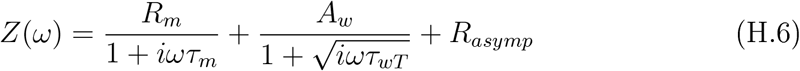

where *τ_wT_* = 1/*ω_wT_*. We calculate 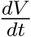 from Equations H.4 and H.5.

Applying Eq. H.5(a) on the right-hand part of Expression H.6 gives [37]).

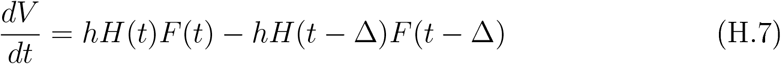

where

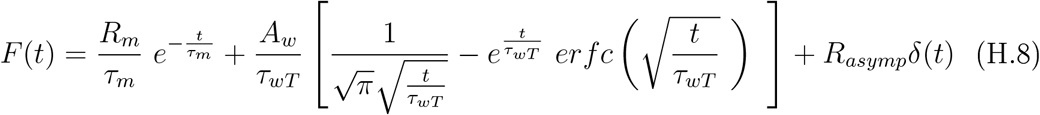

The function 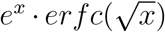*e^x^* for *x* ≥ 0 is a real and positive monotonously decreasing function which tends to zero at infinite. At *x* = 0, it is equal to 1.

Because the experimental measurements shown here indicates that *τ_m_* ≪ *τ_wT_*,we develop F(t) in series around zero to evaluate this function when *t* < *τ_wT_*. We obtain

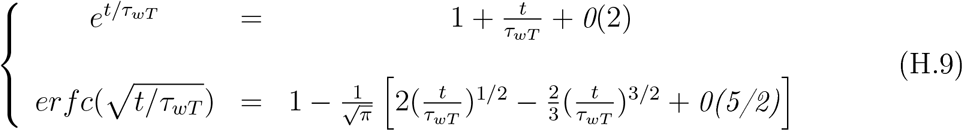

We then write the following equalities:

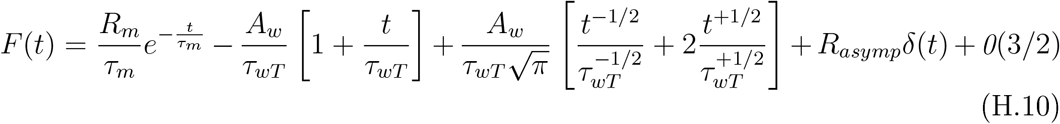

and

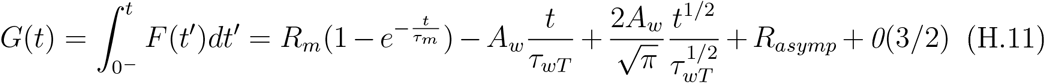

The voltage for *V*(0^−^) = 0 at *t* = 0 and *t* ≥ Δ is given by (H.7)

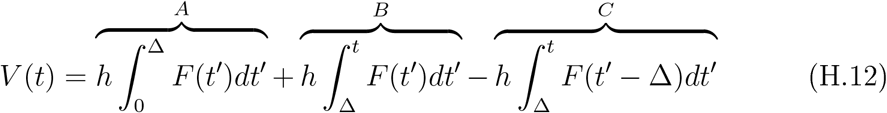

which gives

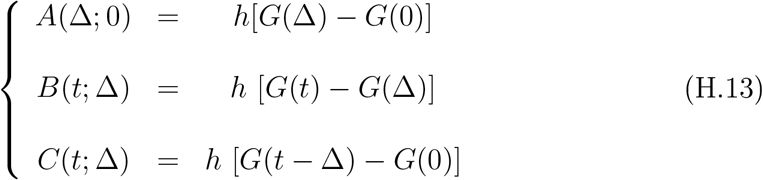

Thus, we have

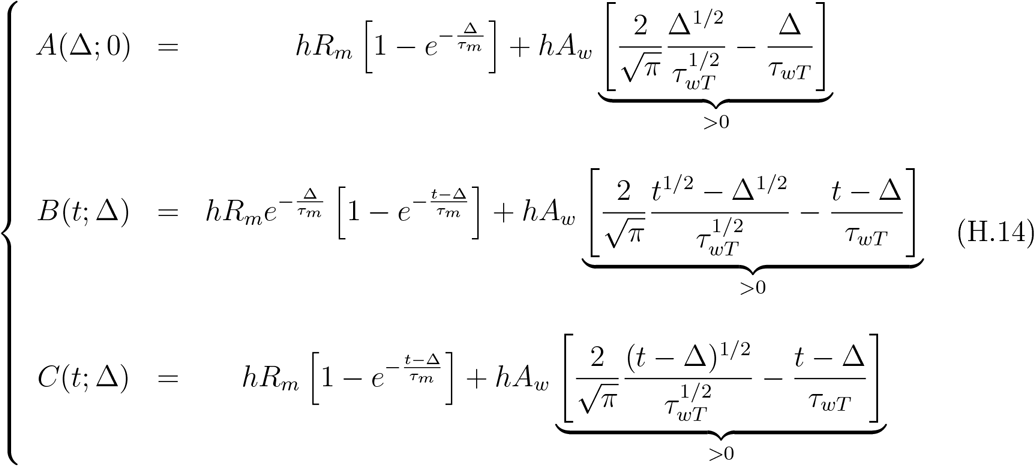

*A*(Δ) corresponds to the voltage reached at time Δ (Δ is the duration of the current pulse). *A*(Δ) depends on *A_ω_*. If *A_ω_* = 0, the voltage is equal to that of a resistive model. In the case of a diffusive model *A_ω_* ≠ 0 is positive because 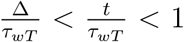. Consequently, the voltage as a function of time is given by:

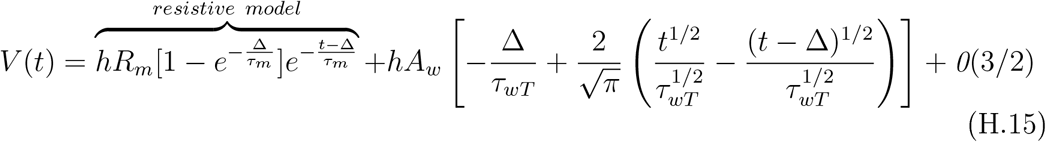

when *t* > Δ and *t* <*τ_wT_*.

Finally, the last expression is equivalent to the second-order approximation in 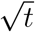 of the expression

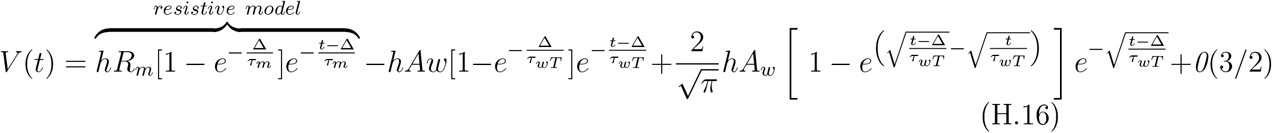

when *t* > Δ and *t* <*τ_wT_*. We can also develop the third term of expression H.16 as a series of exponentials linear in *t* (using Newton binomial) with a combination of exponentials with different relaxation times.

**Figure H.1:**
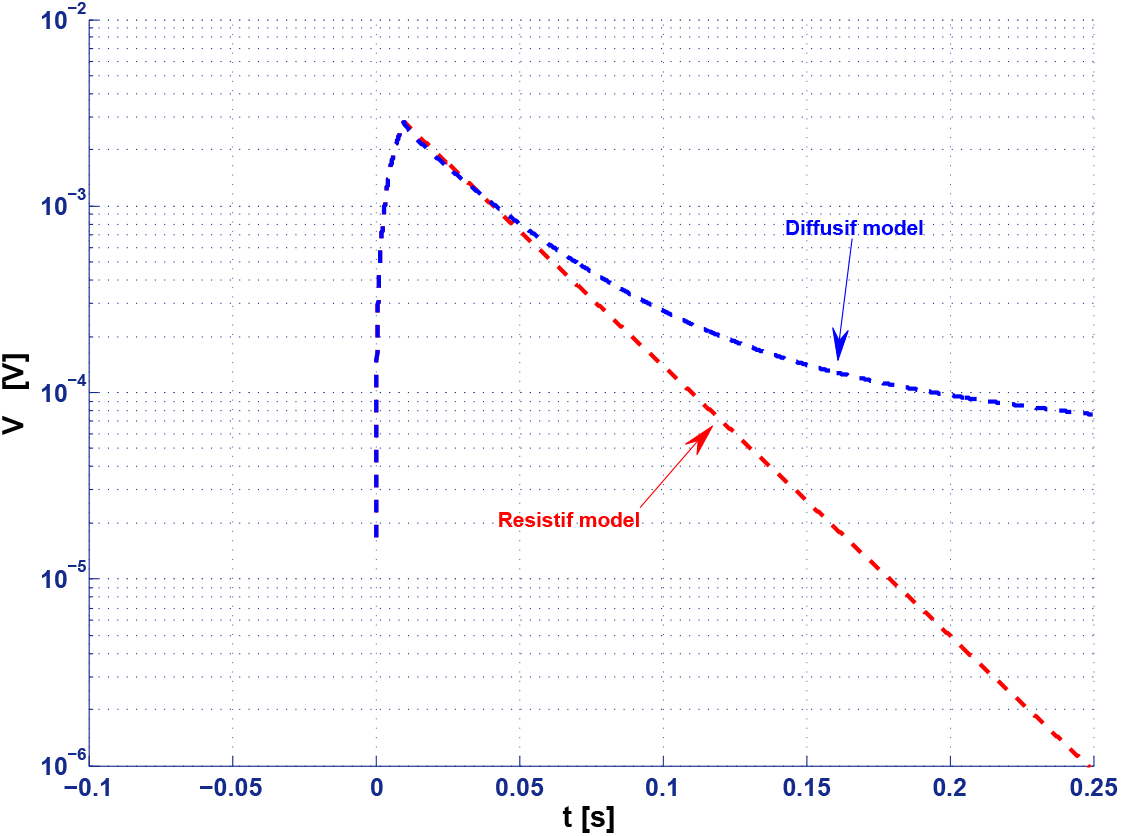
Voltage relative to rest as a function of time during the injection of a current pulse of 10 *pA* and 10 ms duration. The membrane time constant was *τ_m_* = 30 *ms* and the impedance of the extracellular medium + membrane is of 1000 *M*Ω at null frequency. The red curve shows the prediction of the resistive model, while the blue curve shows the resistive model (with a threshold frequency of *ν_wT_* = 0.5 *Hz* and *A_ω_* = *R_m_*).

Consequently, the method of current pulse injection and the linearization method give a different slope in a semi-log graph, according to the type of model. For a resistive model, the slope is −1/*τ_m_*, and this allows to directly estimate the membrane time constant, as classically performed. However, for a diffusive model, the slope is slightly variable and smaller than the resistive model. It depends on *τ_m_*, *ν_wT_* and on the ratio *R_m_*/*A_w_*. For *t* > Δ, one part of the voltage, *V_a_* (*V* = *V_a_* + *V_b_*), attenuates according to the resistive model, while the other part, *V_b_*, attenuates slower and depends on *ν_wT_*. This explains why the membrane time constant seems larger with the pulse or linearization method, compared to the fitting of experimental measurements (Appendix G). The divergence originates from the pulse or linearization methods according to a resistive model.

## Footnotes

1 Thc electric quasistatic approximation consists of neglecting electromagnetic induction such that we have 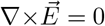. It is important to note that in this approximation, the displacement current is taken into account and accumulation of charges can occur. There exists another approximation, the magnetic quasistatic approximation, where electromagnetic induction is not neglected [21].

2 The term “macroscopic impedance” that we use here corresponds to the definition of the impedance as in electronics, and is different from the macroscopic impedance used at large scales (centimeters).

3 In principle, *F*(*ω*) accounts for other physical phenomena (such as electromagnetic induction, electric viscosity, etc) but these phenomena seem to have a negligible impact on the impedance values for the frequency range < 1000 *Hz* investigated here [7].

4 Wc refer to “quasi-homogeneous” the extracellular medium in culture, because it is very close to a homogeneous medium, and contrasts with the high heterogeneity seen in acute brain slices.

5 Notc that in [10], wc have shown that if there is a strong spatial dependence of the electric parameters, a frequency dependence of the impedance appears. The equation used is equivalent to the generalized current conservation law. The latter can be written as 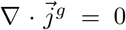 where 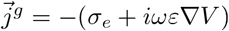, and we have 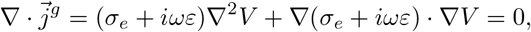 which is equivalent to [10].

6 Note that the diffusive model stems from the application of quasi-static statistical thermodynamics to the model of Gouy-Chapmann-Stcrn-Dcbyc [20, 29].

7 By definition, the external (or internal) Debye layer is the region at the interface between the membrane and the extracellular (or intracellular) medium. It was estimated that the electric potential over this region attenuate by *e* ≈ 2.718 times its value on the membrane surface. Its thickness is called the Debye length. For more details on the notion of Debye layers, sec [29].

8 For example, if we have two resistances in parallel, the majority of the charges will go through the smallest resistance. In electromagnetism, the principle of least constraint can be seen as a generalization of this example [33].

9 Notc that one study (Wilson et al. 2014) reports a frequency-dependent cxtraccllar impedance, but a very different frequency range, between 4.7 kHz and 2 λnĩz. was used.

10 A similar approach is classically used to model the electrical point effect [32].

11 Representing the electric parameters in Fourier frequency space is particularly efficient when the medium is linear because in this case the density of frcc-chargc current and displacement current are proportional to *e^iωt^* if the electric field is also proportional to this term.

## References

[1] Buzsáki, G., Anastassiou, C., and Koch, C. 2012. The origin of extracellular fields and currents: EEG, ECoG, LFP and spikes. Nature Reviews Neurosci. 13, 407–420.

[2] Makarova, J., Gomez-Gala, M., and Herreras, O. (2008). Variations in tissue resistivity and in the extension of activated neuron domains shape the voltage signal during spreading depression in the CAI in vivo. European Journal of Neuroscience 27: 444–456.

[3] Rail, W., G.M. Shepherd. 1968. Theoretical Reconstruction of Field Potentials and Dendrodendritic Synaptic Interactions in Olfactory Bulb J Neurophysiol. Nov;31(6):884–915. doi: 10.1152/jn.1968.31.6.884.

[4] Ranck, J.. 1963. Analysis of specific impedance of rabbit cerebral cortex. Exp. Neurol. 7, 144–152.

[5] Logothetis, N.K., C. Kayser, and A. Oeltermann. 2007. In vivo measurement of cortical impedance spectrum in monkeys: implications for signal propagation. Neuron 55: 809–823.

[6] Miceli, S., T.V. Ness, G.T. Einevoll, and D. Schubert 2017. Impedance Spectrum in Cortical Tissue: Implications for Propagation of LFP Signals on the Microscopic Level. eNeuro, Jan-Feb; 4(1).

[7] Gabriel, S., R.W. Lau, and C. Gabriel. 1996. The dielectric properties of biological tissues: II. Measurements in the frequency range 10 Hz to 20 GHz. Phys. Med. Biol., 41, 2251.

[8] Wagner, T., U. Eden, J. Rushmore, C.J. Russo, L, Dipietro, F. Fregni, S. Simon, S. Rotman, N.B. Pitskel, C. Ramos-Estebanez, A. Pascual-Leone, A.J. Grodzinsky, M. Zahn, and A. Valero-Cabre. 2014. Impact of brain tissue filtering on neurostimulation fields: a modeling study. Neuroimage, Jan 15; 85(0 3): 1048–1057.

[9] Gomes, JM, C. Bédard, S. Valtcheva, MJ. Nelson, V. Khokhlova, P. Pouget, L. Ve-nance, T. Bal and A. Destexhe. 2016. Intracellular impedance measurements reveal non-ohmic properties of the extracellular medium around neurons. Biophys. J. 110: 234–246.

[10] Bédard, C., H. Kröger, A. Destexhe. 2004. Modeling extracellular field potentials and the frequency-filtering properties of extracellular space. Biophys. J. 64:1829–1842.

[11] Bédard, C., H. Kroger and A. Destexhe. 2006. Model of low-pass filtering of local field potentials in brain tissue Phys. Rev. E 73:051911.

[12] Bédard, C., Destexhe, A.. 2009. Macroscopic models of local field potentials and the apparent 1/f noise in brain activity. Biophys. J. 96(7):2589–4608.

[13] Bédard, C., S. Rodrigues, N. Roy, D. Contreras, A. Destexhe. 2010. Evidence for frequency-dependent extracellular impedance from the transfer function between extracellular and intracellular potentials: intracellular-LFP transfer function J. Comput. Neurosci. Dec;29(3):389–403.

[14] Dehghani, N., C. Bédard, S.S. Cash, E. Halgren, A. Destexhe. 2010. Comparative power spectral analysis of simultaneous elecroencephalographic and magnetoencephalo-graphic recordings in humans suggests non-resistive extracellular media. J. Comput. neurosci. 29 (3): 405–421.

[15] Bédard, C., JM. Gomes, T. Bal, A. Destexhe. 2017. A framework to reconcile frequency scaling measurements, from intracellular recordings, local-field potentials, up to EEG and MEG signals. Journal of Integrative Neuroscience 16 (1): 3–18.

[16] Barbour, B. 2017. Analysis of Claims that the Brain Extracellular Impedance Is High and Non-resistive, Biophys. J. Volume 113, Issue 7, 1636–1638.

[17] Bédard, C. and Destexhe, A. 2017. Is the extracellular impedance high and non-resistive in cerebral cortex ?, Biophys. J. Volume 113, Issue 7, 1639–1642.

[18] Nelson, MJ., P. Pouget, E. A. Nilsen, C. D. Patten, and J. D. Schall. 2008. Review of Signal Distorsion through Metal Microelectrode Recordinf Circuits and Filters, J Neurosci Methods, Mar 30; 169(1): 141–157.

[19] Nelson, MJ, S. Valtcheva, L. Venance. 2017. Magnitude and behavior of cross-talk effects in multichannel electrophysiology experiments. J Neurophysiol. 118: 574–594.

[20] Bédard, C., A Destexhe. 2011. Generalized theory for current-source-density analysis in brain tissue, Phys. Rev. E 84 (4):041909.

[21] Le Bellac, M., Lévy-Leblond J.M.. 1973. Galilean electromagnetism. Nuovo Cim B 14, 217–234.

[22] Bédard, C. and Destexhe, A.. 2013. Generalized cable theory for neurons in complex and heterogeneous media. Physical Review E 88:022709.

[23] Rail, W.. 1995. The theoretical foundations of dendritic function. MIT Press, Cambridge, MA.

[24] Tuckwell, H.C.. 1988. Introduction to Theoretical Neurobiology: Linear Cable Theory and Dendritic Structure. Cambridge University Press, Cambdridge, UK.

[25] Bartels, R.H., J.C. Beatty, B.A. Barsky. 1987. An Introduction to Splines for uses in Computer Graphics and Geometric Modeling. Ed. Morgan Kaufman Publishers, Inc, Los Altos, California, 485 p.

[26] Johnston, D. and Wu S.M. 1995. Foundations of Cellular Neurophysiology, Ed. MIT Press.

[27] Maxwell, J.C.. 1873 A Treatise on Electricity and Magnetism. Clarendon Press, Oxford. Vol. 1

[28] Priou, A.. 1992. Dielectric properties of heterogeneous materiels. Progress in Electromagnetics Research, Elsevier, New York.

[29] Vasilicv, A. M.. 1984. Introduction To Statistical Physics, Ed. Mir.

[30] Nicholson, C.. 2005. Factors governing diffusing molecular signals in brain extracellular space. J. Neural Transni. 112, 29–44.

[31] Destcxlie, A. and Bédard C.. 2013. Local Field Potential. Scliolarpedia, http://www.scholarpedia.org/article/Local_field_potential

[32] Kalaclinikov, S.. 1980. Electricity, Ed. Mir.

[33] Goldstein, H., C.P. Poole, J.L. Safko. 2001. Classical Mechanics (3rd Edition). Ed. Addison Wesley, 647 p.

[34] Foster, K. R., H.P. Scliwan. 1989. Dielectric properties of tissues and biological materials: A critical review. Critical Reviews in Biomedical Engineering 17(1):25–104.

[35] Bédard, C., Destcxlie A.. 2014. Mean-field formulation of Maxwell equations to model electrically inhomogeneous and isotropic media Equations to Model Electrically. Journal of Electromagnetic Analysis and Applications 6 (10), 296.

[36] Bédard, C., Destcxlie A.. 2014. Generalized cable formalism to calculate the magnetic field of single neurons and neuronal populations. Physical Review E 90 (4):042723.

[37] Obcrliettinger, F. and L. Badii. 1973. Tables of Laplace Transforms. Springer-Vcrlag Berlin Heidelberg, New York.

[38] Rail, W.. 1995. The Theoretical Foundation of Dendritic Function. Scgcv, L, J. Rinzcl and G.M. Shepherd, ed. MIT Press, Cambridge MA.

[39] Hines, M.L., and N.T. Carnevale. 1997. The NEURON simulation environment. Neural Computation 9: 1179–1209.

[40] Nelson, M.J., C. Bosch, L. Venance, and P. Pougct. 2013. Microscale Inhomogeneity of Brain Tissue Distorts Electrical Signal Propagation. The Journal of Nour oscience, 33 (7):2821–2827.

[41] Hrabetova S, Cognet L, Rusakov DA, Nagerl UV. 2018. Unveiling the extracellular space of the brain: from super-resolved microstructure to in vivo function. J Neurosci. 38: 9355–9363.

[42] Herreras, O. 2016, Local Field Potentials: Myths and Misunderstandings. Front. Neural Circuits, 10: 101. https://doi.org/10.3389/fncir.2016.00101

